# Trisomy 21 Impairs Synchronized Activity in Human Down Syndrome Cortical Excitatory Neuron Networks

**DOI:** 10.1101/2024.12.27.630530

**Authors:** Manuel Peter, Raquel Real, Alessio Strano, Hugh P. C. Robinson, Mark A. Smith, Samuel J. Barnes, Tania Dutta, Vincenzo De Paola, Frederick J. Livesey

## Abstract

Down syndrome (DS) is the most common genetic cause of intellectual disability, affecting one in 700 live births worldwide, and is caused by trisomy of the human chromosome 21 (Hsa21), yet the cellular and molecular mechanisms driving neuronal network dysfunction remain poorly understood. Here, we investigated whether trisomy 21 (TS21) alters the spontaneous synchronised activity of excitatory neuron networks, potentially contributing to the neurodevelopmental phenotypes of DS. By modelling human cortical development *in vitro* with TS21 and matched control human iPSCs from two DS donors, we investigated the impact of Hsa21 triplication on neural network activity and connectivity. Calcium imaging revealed an early and pronounced reduction in neuronal activity in TS21 cortical neurons, including a marked loss of synchronised bursting. These deficits persisted up to 80 days *in vitro* and for over 5 months *in vivo* following transplantation into the mouse forebrain, as shown by multiphoton calcium imaging through a cranial window. Viral trans-synaptic tracing identified significant reduction of neuronal connectivity in TS21 neuronal networks *in vitro*, suggesting that reduced network connectivity contributes to the dramatic reduction of synchronised bursting. Furthermore, TS21 neurons displayed significantly reduced expression of voltage-gated potassium channels, with single-neuron recordings confirming a reduction of hyperpolarization-activated currents. Together, these findings demonstrate long-lasting impairments in human cortical excitatory neuron network function associated with Trisomy 21.

## Introduction

DS is the most common chromosomal abnormality in humans, associated with a number of neurodevelopmental and neurological features, including microcephaly and intellectual disability (Antonarakis et al 2004). TS21 alters the expression of > 200 Hsa21 protein coding genes (Hattori et al 2000), resulting in a gene dosage imbalance believed to underlie several of the phenotypes observed in DS (Asim et al 2015, Wiseman et al 2015). Trisomy 21 in humans commonly results in a range of developmental and morphological changes in the forebrain, including a decrease in neural progenitor cells (Guidi et al 2008), altered cortical lamination (Golden & Hyman 1994) and hippocampus morphology, a reduction of dendritic spines (Ferrer & Gullotta 1990, Suetsugu & Mehraein 1980), and overall reduced brain volume (Pinter et al 2001). In adulthood, individuals with DS face a high-risk of early-onset Alzheimer’s disease, partly due to the APP gene residing on Hsa21, leading to a functional triplication of the APP locus (Wiseman et al 2009). Functionally, altered neuronal activity (Karrer et al 1998, Velikova et al 2011) and synchronization (Babiloni et al 2010) are observed in the brains of individuals with DS, reflecting an altered network architecture characterised by increased local connectivity (Anderson et al 2013, Pujol et al 2015). However, the developmental origins and cellular mechanisms underlying the changes in synchronised brain activity in DS are currently not well understood.

Several mouse models for human TS21 have been developed, which use either syntenic regions of mouse chromosomes or parts of human chromosome 21, many of which partially replicate some aspects of human DS neuropathology (Best et al 2012, Chakrabarti et al 2007, Cooper et al 2012, Cramer et al 2015, Herault et al 2017, Stern et al 2015). Human induced pluripotent stem cells (hiPSCs) and their directed differentiation to model the development of different regions of the CNS provide alternative systems to study human-specific aspects of neurodevelopmental disorders (Brennand et al 2011, Chailangkarn et al 2016). Several TS21 hiPSC lines have been generated (Alonzo et al 2023, Giffin-Rao et al 2022) (Kawatani et al 2021, Park et al 2008) (Weick et al 2013) (Shi et al 2012b) and used to investigate various features of AD pathology and development (Russo et al 2024). In our previous work, DS donor hiPSC-derived neurons showed normal *in vitro* single cell electrophysiological properties, but increased production of Aβ peptides, increased amyloid plaques and mislocation of phosphorylated tau, reminiscent of AD pathology (Shi et al 2012b). We also showed reductions in developing cortical neuron network activity *in vivo* (Real et al 2018). However, in another study, TS21 human neurons exhibited reduced synaptic activity, despite maintaining normal excitability (Weick et al 2013). Furthermore, it remains unclear whether and how synchronized network activity, which is essential for the formation and refinement of functional networks (Martini et al 2021), is affected. Specifically, it is not yet clear whether the reduced activity of pyramidal neurons is an intrinsic property of the excitatory neuron network itself or is influenced by hyperactive interneurons (Zorrilla de San Martin et al 2020) and/or astrocytes (Mizuno et al 2018).

Here, we differentiated isogenic and non-isogenic TS21 and control hiPSCs into predominantly cortical excitatory neurons to examine cell-autonomous effects of the extra Hsa21 copy on neuronal network activity and connectivity during early development. We identified impaired TS21 neuronal network activity development *in vitro* and *in vivo* after transplantation into the somatosensory cortex of mice, with changes in the proportions of deep to upper layer cortical neurons present. We furthermore found that TS21 excitatory neurons were severely under-connected compared to control neurons. Using gene expression analysis and electrophysiology, we discovered a downregulation of multiple voltage gated potassium channels and the hyperpolarization-activated cyclic nucleotide–gated channel HCN1 in TS21 neurons. Collectively, the observed reductions in ion channel expression, neuronal composition, neuronal connectivity, and network activity synchronization may contribute to functional differences relevant to the cognitive and intellectual features associated with DS.

## Results

### Differentiation of TS21 and control hiPSCs to cortical excitatory neurons

We differentiated two TS21, DS hiPSC lines TS21-1, (DSIPS), (Park et al 2008), TS21-2, (Tho3B), (Real et al 2018) and two WT, control lines WT-1, (NDC1.2), (Israel et al 2012), WT-2, (Tho3F), (Real et al 2018) towards a cortical forebrain identity as previously described (Shi et al 2012a). Notably, TS21-2 and WT-2 are an isogenic pair, from the same donor, where the third copy of Hsa21 was lost in WT-2 during re-programming (Real et al 2018). Immunofluorescence staining for the neural progenitor marker Pax6 and the proliferation marker Ki67 confirmed robust neuronal induction and abundant mitotically active neural progenitor cells (Figure 1A) that gave rise to Tbr1, Ctip2 (Figure 1B), and Tuj1 (Figure 1C, Figure 1—figure supplement 1) deep-layer cortical neurons at day 50 in both control and TS21 lines.

**Figure 1.**
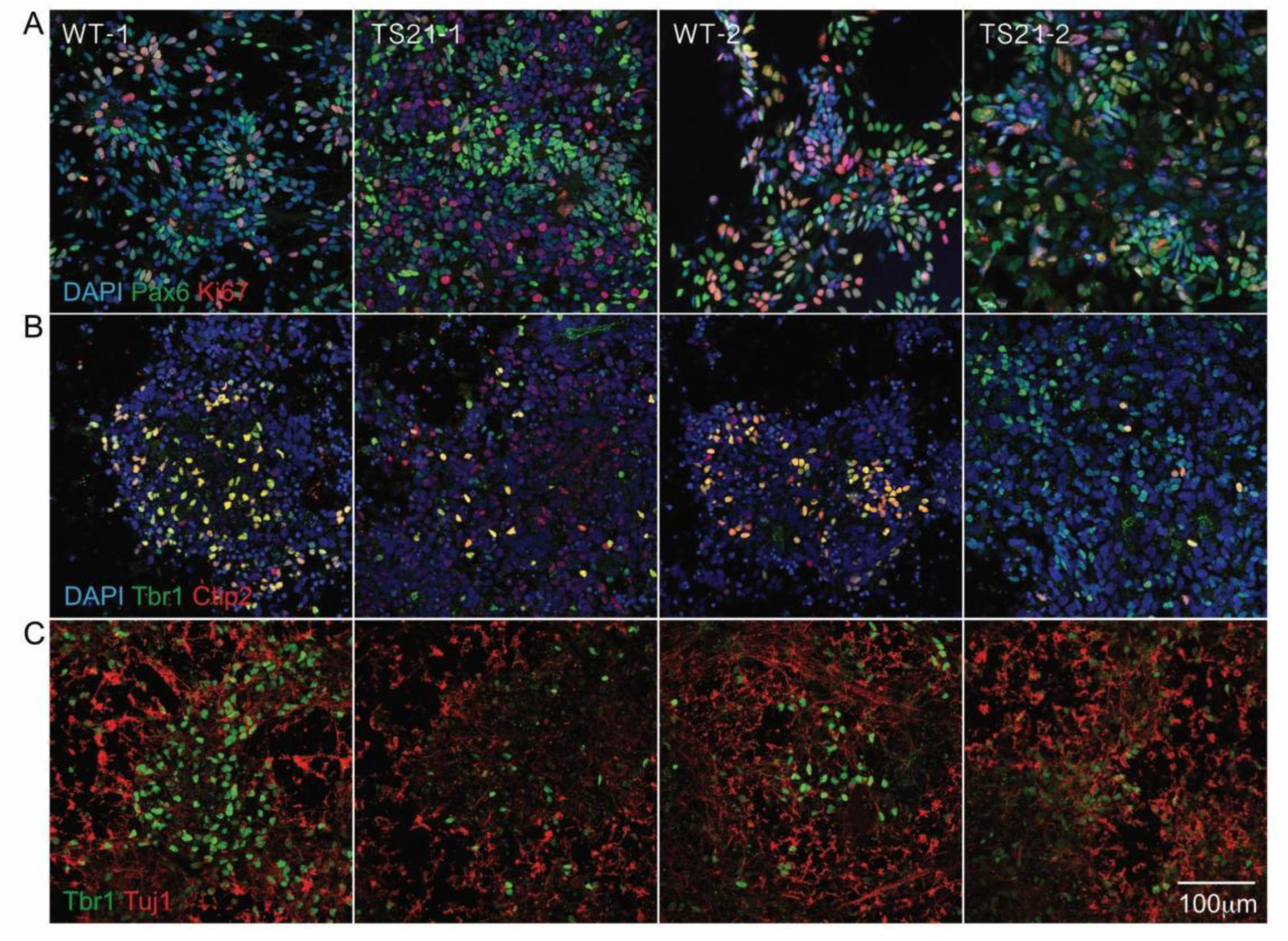
*In vitro* differentiation of control (WT) and trisomic (TS21) iPSCs into neural progenitor cells and cortical excitatory neurons. A: Confocal image stacks of neural progenitor cells of two control (WT-1, WT-2) and two trisomic (TS21-1, TS21-2) cultures at day 35 – 37. B: Confocal image stacks of WT and TS21 trisomic neurons at day 50, immunofluorescence staining for the deep layer marker Tbr1 and Ctip2. C: Confocal image stacks of WT and TS21 neurons at day 50 immunofluorescence staining for the deep layer marker Tbr1 and the pan neuronal marker Tuj1.

Whole cell electrophysiological recordings on day 53/54 showed a large proportion of TS21 and control neurons firing action potentials after current injection (Figure 1—figure supplement 2A, D). We did not detect significant differences in resting membrane potential (WT-1: n = 20, –36±14mV; TS21-1: n=21, –31±14mV; WT-2: n=22, –32±13mV; TS21-2: n=19, –27±8mV; mean±SD, Fig 1B), evoked action potential amplitude (WT-1: n=14, 14±10mV; TS21-1: n=18, 16±11mV; WT-2: n=17, 19±13mV; TS21-2: n=13, 15±11mV; mean±SD, Figure 1—figure supplement 2C) or in voltage gated Na^+^ currents (Figure 1—figure supplement 2E), indicating no differences in whole cell properties at this early developmental timepoint.

### Excitatory neural network activity development is impaired in DS neurons

Human cortical neural networks undergo stereotypical activity changes during development, with an early phase of uncorrelated firing, followed by a phase of highly synchronised bursting, and lastly the emergence of highly structured, complex firing patterns later in development (Kirwan et al 2015). These complex patterns are similar to synchronised bursting *in vivo* (Khazipov et al 2004, Kilb et al 2011, Leinekugel et al 2002) and in cortical brain slices (Corlew et al 2004), and are believed to play a role in the maturation of neuronal circuits.

To test if synchronised bursting also occurs in TS21 cortical excitatory networks *in vitro*, we loaded cortical excitatory neurons derived from the two WT (WT-1, WT-2) and two TS21 hiPSC lines (TS21-1, TS21-2) with the membrane permeable calcium sensitive dye Oregon Green 488 BAPTA-1, AM (OGB), and recorded fluorescence changes over two minutes (Figure 2). Changes in the fluorescence signal (ΔF/F) served as a readout for neuronal activity and allowed the simultaneous recording of the neuronal activity of hundreds of neurons at single-cell resolution. As previously reported (Kirwan et al 2015), we found that both WT lines acquired spontaneous synchronized bursting activity between day 50 and 60 (Figure 2—figure supplement 1B), which was sensitive to the reversible effect of the Na^+^ channel blocker tetrodotoxin (TTX) (Figure 2—figure supplement 1C, left).

**Figure 2.**
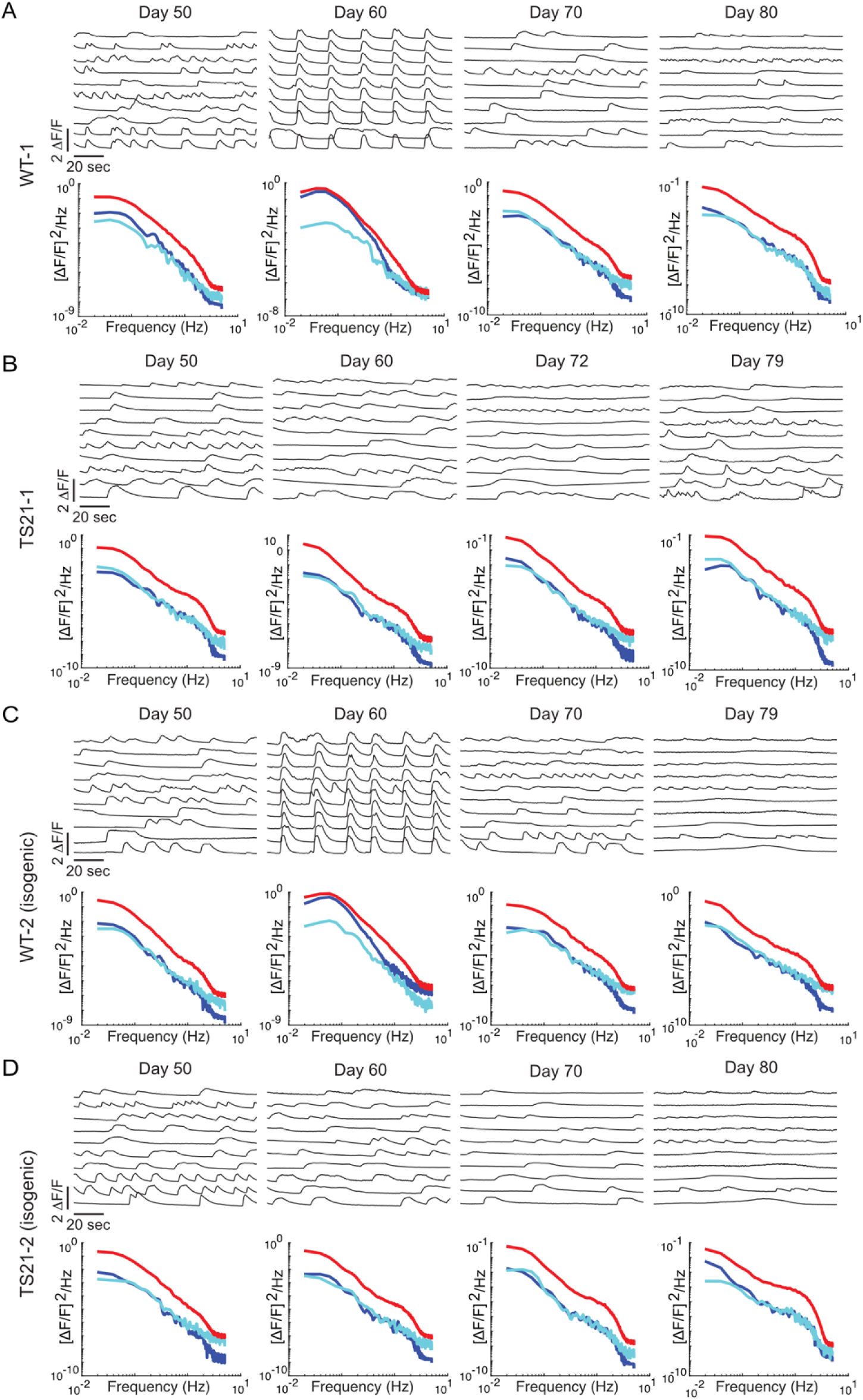
Altered spontaneous network activity development in TS21 neuronal networks *in vitro*. A, B, C, D: Top: Traces of fluorescence changes (ΔF/F) in WT (WT1-1, WT-2) and trisomic (TS21-1, TS21-2) neurons loaded with the calcium indicator OGB at different developmental stages. Each trace shows a 2 minutes time series of the activity of a single neuron (the 10 most active neurons per ROI are shown). Note the absence of synchronized activity at day 60 in both trisomic lines compared to WT control lines. Importantly, synchronized network activity is reinstated in the isogenic, disomic control line WT-2. Bottom: Corresponding power spectra of the 50 most active neurons, red line: average power of all neurons, blue line: power spectrum of the network, cyan line: randomized network power. Note that for both WT networks at day 60 the power spectrum of the network is similar to the average power of all neurons indicating network synchronization.

Changes in network synchronicity were quantified by calculating power spectra for the average network activity (blue) and the average activity of individual neurons (red) (Figure 2—figure supplement 1C, D, E). In a synchronized network, the network power spectrum for a given frequency is close to the power spectrum of a single neuron. In unsynchronized networks the network power spectrum will be close to its own randomized power spectrum (cyan). Control synchronized networks had network power spectra overlapping with the power spectrum of individual neurons at 0.1Hz, indicating a high degree of synchronization at this frequency. Voltage-gated sodium channel blockade with tetrodotoxin (TTX) led to a complete loss of synchronized activity and the network power spectrum overlapped with its own, randomized power spectrum. After removal of TTX, neuronal firing returned to levels similar to before TTX application and showed a high degree of synchronicity (Figure 2—figure supplement 1C, right). Glutamate receptors are important mediators of excitatory network activity. Therefore, we tested if synchronized neuronal network activity is α-amino-3-hydroxy-5-methyl-4-isoxazolepropionic acid (AMPA) or N-methyl-D-aspartate (NMDA) receptor-dependent (Kirwan et al 2015, Kuijlaars et al 2016) by recording neuronal activity of WT neurons in the presence of the AMPA receptor antagonist CNQX (Figure 2—figure supplement 1D) or the NMDA receptor antagonist AP5 (Figure 2—figure supplement 1E). Both drugs led to a complete and reversible loss of synchronicity, although non-synchronized, spontaneous single neuron firing was still present under both conditions. We further noted a small decrease in the average neuronal activity, as indicated by a decrease in the power spectrum (red). Taken together, these results replicate previous findings (Kirwan et al 2015), further demonstrating that spontaneous synchronized network activity is caused by excitatory network activity and is AMPA and NMDA receptor dependent.

To test if synchronized bursting behavior also develops in TS21 neuronal networks, we compared spontaneous network activity of TS21 and WT control neurons over a 30 day period. As reported previously (Kirwan et al 2015), WT neurons showed strong but uncorrelated or weakly synchronized activity at day 50. By day 60, most of the WT neurons exhibited a high degree of synchronized bursting, which disappeared by day 70 *in vitro* (Figure 2A, C). Importantly, a phase of synchronized bursting was never detected in TS21 neurons, with no evidence for synchronized firing in TS21 networks up to day 80 (Figure 2B, D), the last point at which we assayed activity. The loss of synchronized activity was caused by the extra copy of Hsa21, as isogenic control (WT-2) networks developed synchronized activity similar to the independent control line (WT-1) (Figure 2A, C).

We further calculated the mean integral of the power spectra over 0.1 and 1 Hz and noticed a small but significant decrease of averaged neuronal network activity in both TS21 lines (TS21-1: n=17, 0.025±0.009; TS21-2: n=24, 0.016±0.014), compared to WT neurons at day 50 (WT-1: n=22, 0.112±0.082; WT-2: n=25, 0.131±0.103; WT-1 vs. WT-2: ns, WT-1 vs. TS21-1: **, WT-1 vs. TS21-2: ***, WT-2 vs. TS21-1: ***, WT-2 vs. TS21-2: ***, TS21-1 vs. TS21-2: ns, Tukey’s multiple comparisons test, *P <0.05) (Figure 2—figure supplement 2A, B).

WT cells consistently exhibited higher amplitudes and frequencies than TS21 cells, but differences did not reach statistical significance due to variability and limited sample size (Figure 2—figure supplement 3A, B). Nonetheless, effect sizes (Cohen’s d) were large, indicating a strong underlying biological difference in network activity between WT and TS21 neurons. Taken together, these results demonstrate that TS21 neurons do not undergo normal network activity development and lack a phase of spontaneous synchronized bursting activity present in WT neurons.

### Transplantation and *in vivo* two-photon Ca2+ imaging confirms synchronization deficits in TS21 excitatory neural networks

Because the *in vitro* networks were intentionally designed to be predominantly excitatory, with only a small number of interneurons emerging at later stages and no integration into broader CNS circuitry, they provide a restricted view of neuronal maturation and function. By contrast, transplantation into the mouse forebrain exposes control and TS21 neurons to a physiologically relevant milieu that includes vascularization, immune cells, and host-derived signals. Although the transplanted human cells do not fully integrate into host circuits (Real et al 2018), they still achieve greater maturation and more balanced network activity than is possible *in vitro* (Revah et al 2022), allowing us to longitudinally monitor their activity under conditions that better approximate the *in vivo* brain environment. To specifically record from the engrafted neurons, we transduced neural progenitor cells and immature neurons from each genotype with a lentiviral vector expressing the red fluorescence protein tdTomato and the genetically-encoded calcium sensor GCaMP6s (Chen et al 2013). We then transplanted this mixture of neural progenitor cells and immature (day 35-38) neurons into the somatosensory cortex of immunodeficient adult mice (same cohort as in (Real et al 2018)) and implanted a small cranial window, for *in vivo* imaging, on top of the cortex (Real et al 2018) (Figure 3A). Cortical identity of transplanted neurons was confirmed by expression of the transcription factor Tbr1 (expressed in Layer 6 projection neurons) and the human-specific nuclear marker hNu at the same time points where we studied network activity using *in vivo* calcium imaging (i.e. 5 to 6 months post transplantation) (Figure 3B). Notably, the expression of SATB2, a marker for upper-layer cortical projection neurons, was significantly reduced in TS21 grafts at the same time points (Figure 3—figure supplement 1). The observed reduced generation of upper layer neurons from TS21 iPSC-derived neurons is consistent with previous reports from human post-mortem material (Klein & Haydar 2022, Russo et al 2024).

**Figure 3.**
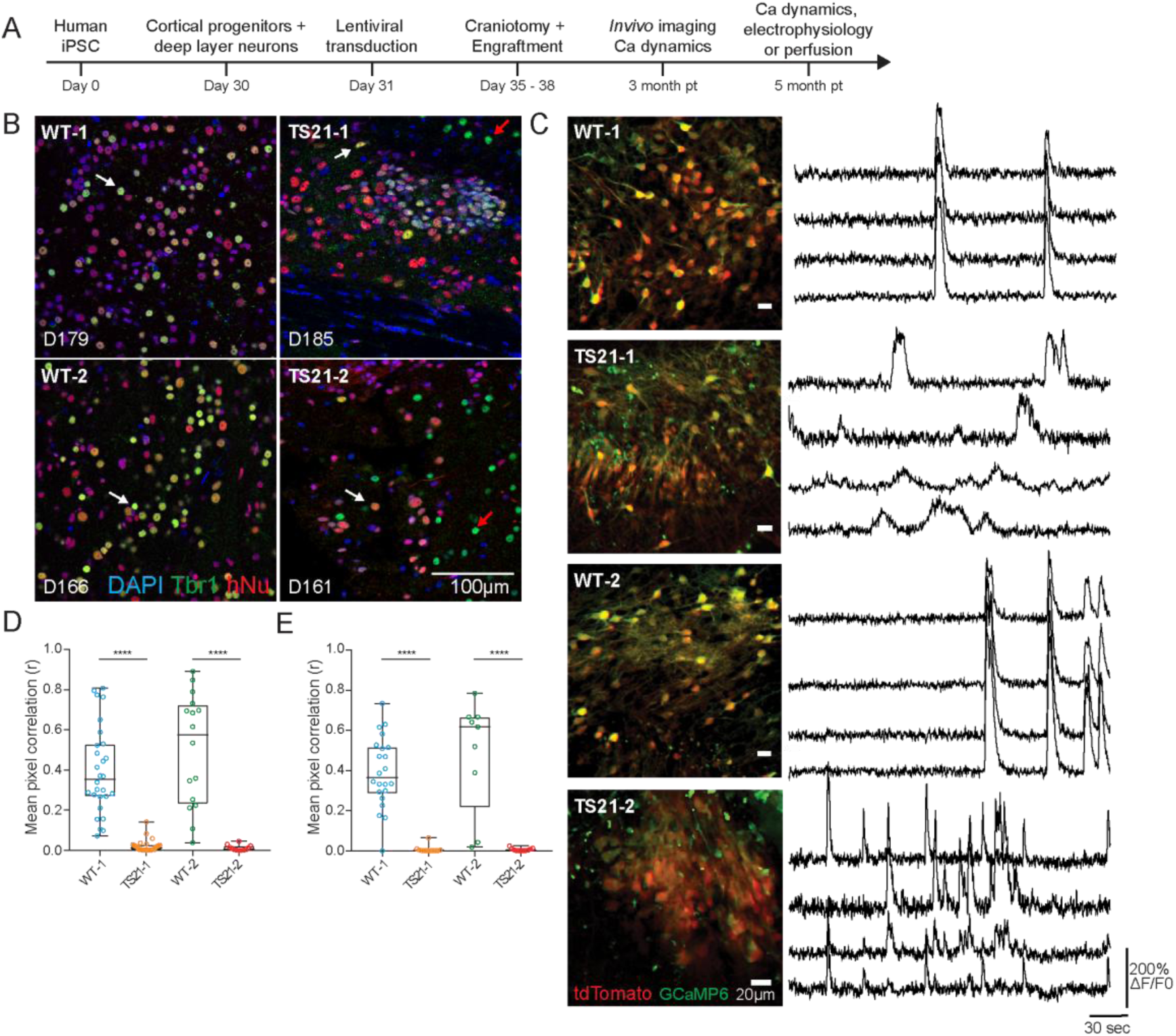
Altered network activity in trisomic (TS21) grafted neurons. A: Schematic of experimental design. B: iPSC derived human cortical neurons transplanted into the somatosensory cortex of mice. Representative confocal images of a graft section immunostained for the deep layer marker Tbr1 and the human specific nuclear marker (hNu). White arrows indicate human neurons double positive for Tbr1 and hNu. Red arrows indicate Tbr1 positive mouse neurons. Days post-transplantation are indicated. C: Left: example images of a two-photon time-series of tdTomato and GCaMP6 expressing neurons 3 months post transplantation. Right: Traces of fluorescence changes (ΔF/F0) in control and trisomic neurons. Note the absence of synchronized activity in both cell lines of TS21 neurons compared to WT neurons. D: Mean pixel correlation (r) measured 3 months after graft implantation for WT-1 (n = 28 ROIs), TS21-1 (n = 26 ROIs), WT-2 (n = 16 ROIs) and TS21-2 (n = 12 ROIs) grafts. Mann-Whitney U-test, ****P < 0.0001. E: Mean pixel correlation (r) measured 5 months after graft implantation for WT-1 (n = 22 ROIs), TS21-1 (n = 12 ROIs), WT-2 (n = 9 ROIs) and TS21-2 (n = 11 ROIs) grafts. At least 3 mice per line per condition (3 mpt and 5 mpt). Mann-Whitney U-test, ****P < 0.0001. Median, interquartile range, minimum and maximum values are indicated. Each data point represents an ROI.

To confirm that the transplanted neurons were functional, we performed patch-clamp recordings in acute brain slices 5 months after transplantation. Grafted cells were identified based on fluorescence and selected for patching based on the presence of pyramidal cell morphology. We did not detect significant changes in the resting membrane potential (WT-1: n=18 cells, –54±7 mV; TS21-1: n=21 cells, –49±13 mV; WT-2: n=27 cells, –57±9 mV; TS21-2: n=16 cells, –55±11 mV, mean±SD, Figure 3—figure supplement 2A) or the input resistance (WT-1: n=18 cells, 1.34±0.63 GΩ; TS21-1: n=21 cells, 1.83±1.20 GΩ; WT-2: n=27 cells, 0.97±0.75 GΩ; TS21-2: n=16 cells, 1.35±1.07 GΩ, mean±SD, Figure 3—figure supplement 2B) between all four lines. However, a significant decrease in membrane capacitance was present when comparing disomic WT-1 with trisomic TS21-1 neurons (WT-11: n=18 cells, 19.44±9.41 pF; TS21-1: n=21 cells, 13.06±7.07 pF; WT-2: n=27 cells, 24.58±11.96 pF; TS21-2: n=16 cells, 20.9±13.2 pF, mean±SD (Figure 3—figure supplement 2C).

Neurons from all four lines fired action potentials after current injection (Figure 3—figure supplement 2D-I). The frequency of evoked action potentials upon incremental steps of current injection was similar between WT-1 vs TS21-1 and WT-2 vs TS21-2, with the exception of at +50 pA between WT-2 vs TS21-2 (Tukey’s multiple comparisons test after two-way ANOVA, interaction *F*_12,321_ = 0.3128, *P* = 0.9869; *\*P* < 0.05; *ns*, not significant, Figure 3—figure supplement 2L). Evoked action potential amplitudes (WT-1: n=18 cells, 91.3±11.1 mV; TS21-1: n=14 cells, 77.0±16.4 mV; WT-2: n=24 cells, 87.9±16.0 mV; TS21-2: n=10 cells, 82.7±10.8 mV, mean±SD, Figure 3—figure supplement 2J) and widths (WT-1: n=18 cells, 2.20±0.74 ms; TS21-1: n=16 cells, 1.59±0.48 ms; WT-2: n=24 cells, 2.13±0.99 ms; TS21-2: n=10 cells, 1.48±0.42 ms, mean±SD, Figure 3—figure supplement 2K) were slightly reduced in TS21 neurons, which could be attributed to differences in neuronal maturation or electrogenesis between cells from the two genotypes.

Finally, we recorded spontaneous network activity in the grafts by *in vivo* imaging of GCaMP6-expressing neurons (Figure 3C). While neurons in all grafts had spontaneous baseline neural activity at 3 months post-transplantation, we found a significant reduction in mean pixel-wise correlation across entire ROIs in both TS21 lines (Figure 3D), indicating decreased synchronized neuronal activity between multiple somas and neuropil. To exclude that this synchronization deficit was not due to a delay in functional maturation of TS21 networks, we repeated recordings of spontaneous activity 5 months after transplantation (Figure 3E). Again, we observed a strong synchronization defect in TS21 neurons.

This deficit in network synchronization was associated with the presence of an extra copy of Hs21 and was reversed by normalization of the Hsa21 copy number in the isogenic control (WT-2 cell line).

### Reduced connectivity in TS21 neuronal networks

To test whether changes in connectivity could underlie the synchronized network activity defect present in TS21 neural networks, we assayed the connectivity of DS and control neurons using a pseudotyped, monosynaptic spreading rabies virus (Brennand et al 2011, Wickersham et al 2007). To do so, we transduced neurons with low titer lentivirus (LV-SYN-HTG) expressing a histone H2B-GFP fusion, the TVA receptor and a glycoprotein (H2B-GFP::TVA::G), under the control of the human synapsin 1 promoter. After 10 days, we transduced neurons expressing TVA with a pseudotyped mCherry-expressing rabies (Rabies-mCherry) virus. Since high levels of rabies virus can have negative effects on cell survival, we limited rabies spread to 7 days. Primary infected neurons were double positive for H2B-GFP and mCherry. Monosynaptically connected neurons to those primary infected neurons were positive for mCherry only (Figure 4C).

**Figure 4.**
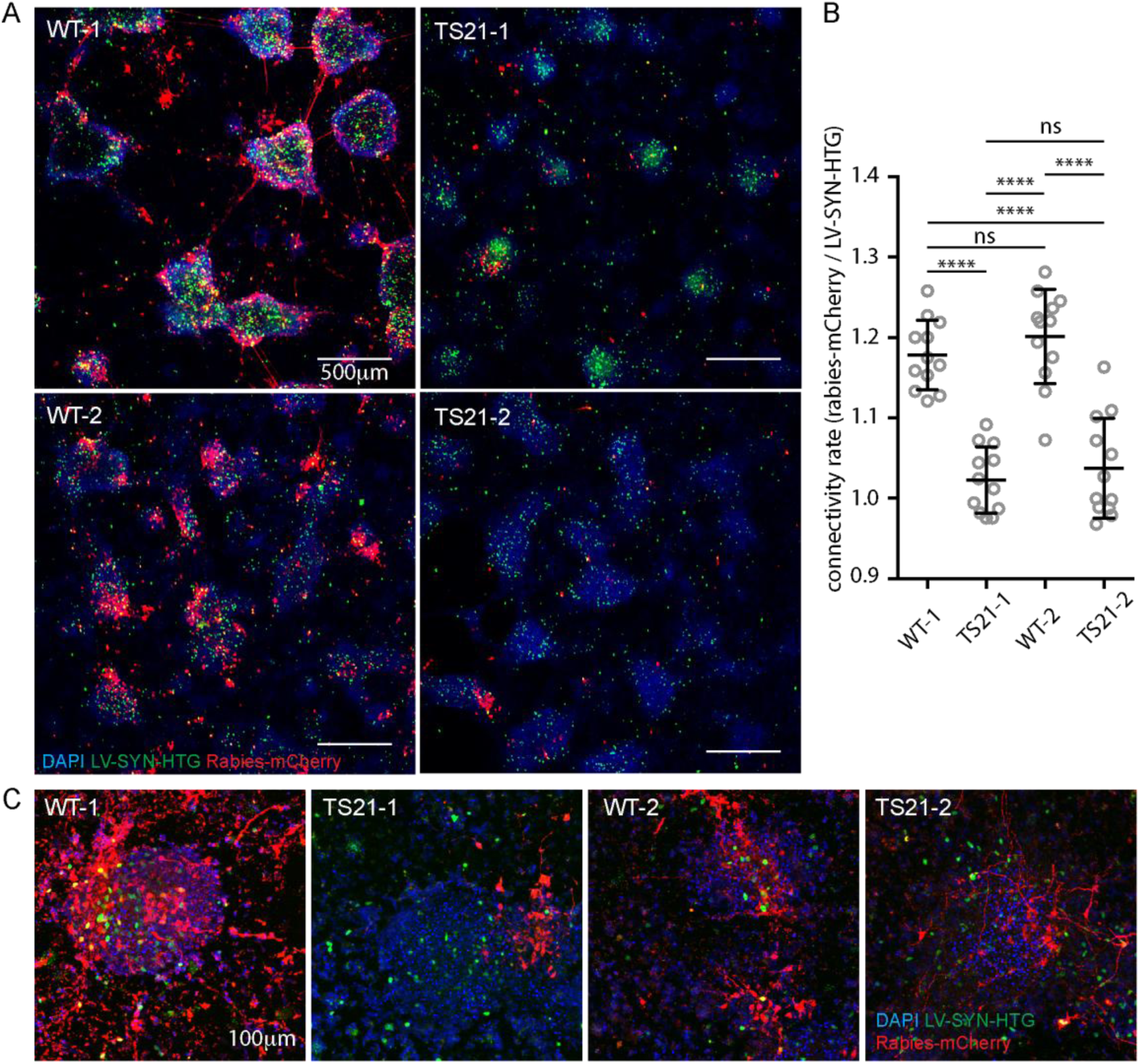
Reduced connectivity in trisomic (TS21) neurons. A: Representative images of WT and trisomic neurons, co-infected with a lentivirus expressing TVA and nuclear GFP (TVA-NLS-GFP) and a pseudotyped mCherry expressing rabies virus (Rabies-mCherry). B: Graph showing the relative red to green pixel values of WT and trisomic neuronal cultures. Note the significant decrease in TS21 neurons compared to WT neurons indicating a lower level of connectivity (WT-1: 1.18 ± 0.04; TS21-1: 1.02 ± 0.04; WT-2: 1.20 ± 0.06; TS21-2: 1.04 ± 0.06, one-way ANOVA, n = 12 ROIs from 4 wells of one neuronal induction per genotype). C: Confocal image stacks of neurons in A taken at higher resolution. Note that connected neuronal clusters (red neurons connected to the ones with green nuclei) can be found in WT and Ts21 neurons.

To score connectivity, we used an automated high-throughput imaging system to take images of the middle of the culture dish in an unbiased way (Figure 4A), scoring average connectivity as the number of mCherry positive neurons relative to GFP positive neurons. We calculated a connectivity value consisting of the ratio of the red to green fluorescence pixel values, with higher values indicating higher connectivity (Brennand et al 2011). This approach normalised for differences in primary infection levels of the lenti– and rabies virus. Using this metric, there was a significant decrease in connectivity among TS21 neurons compared to control neurons (WT-1: 1.18 ± 0.04; TS21-1: 1.02 ± 0.04; WT-2: 1.20 ± 0.06; TS21-2: 1.04 ± 0.06; mean ± SD; n = 4 wells) (Figure 4B). This connectivity defect was specific to TS21 neurons as it was not present in disomic isogenic control neurons (WT-2), which had connectivity rates indistinguishable from the unrelated euploid (disomic) control line (WT-1). Furthermore, the observed decrease in connectivity in TS21 neurons was not due to the inability of the rabies virus to transfer between TS21 neurons, as we were able to detect clusters of mCherry-only positive neurons in all four lines, but to a much lower extent in TS21 neuronal cultures (Figure 4C).

The mechanism underlying retrograde monosynaptic rabies spread is not fully understood and it is conceivable that virus spreading is influenced by neuronal activity. In that case, our observed decrease in connectivity in trisomic neurons could be explained by their lower basal network activity. We tested this hypothesis by adding the sodium channel blocker TTX to the culture medium 24h after rabies virus infection and for the 7 days thereafter. We did not detect a significant difference in connectivity rates between TTX (0.88 ± 0.02) and mock treated neurons (0.85 ± 0.02), indicating no effect of neuronal activity on rabies virus spreading between neurons (Figure 4—figure supplement 1). Taken together, these results indicate that the extra copy of Hsa21 reduces neuronal connectivity among TS21 neurons.

### Voltage gated potassium (K+) channel expression is altered in TS21 neurons

Ionotropic glutamate receptors mediate synaptic transmission, plasticity and memory function (Rao & Finkbeiner 2007). Several DS mouse models show deficits in hippocampal dependent learning, as well as LTP and LTD induction, indicating a role of AMPA and NMDA receptors in DS (Kleschevnikov et al 2004, O’Doherty et al 2005, Siarey et al 1999, Siarey et al 2005). Since we found that spontaneous, synchronised network activity is AMPA and NMDA receptor dependent (Figure 2—figure supplement 1D, E), we asked if the lack of synchronicity in TS21 neurons could be caused by dysregulation or loss of ionotropic glutamate receptors. We performed an RNAseq analysis on day 50 and found 235 genes that were significantly upregulated and 160 genes that were significantly downregulated in trisomic versus WT control neurons (Figure 5A, Figure 5—figure supplement 1-2).

**Figure 5.**
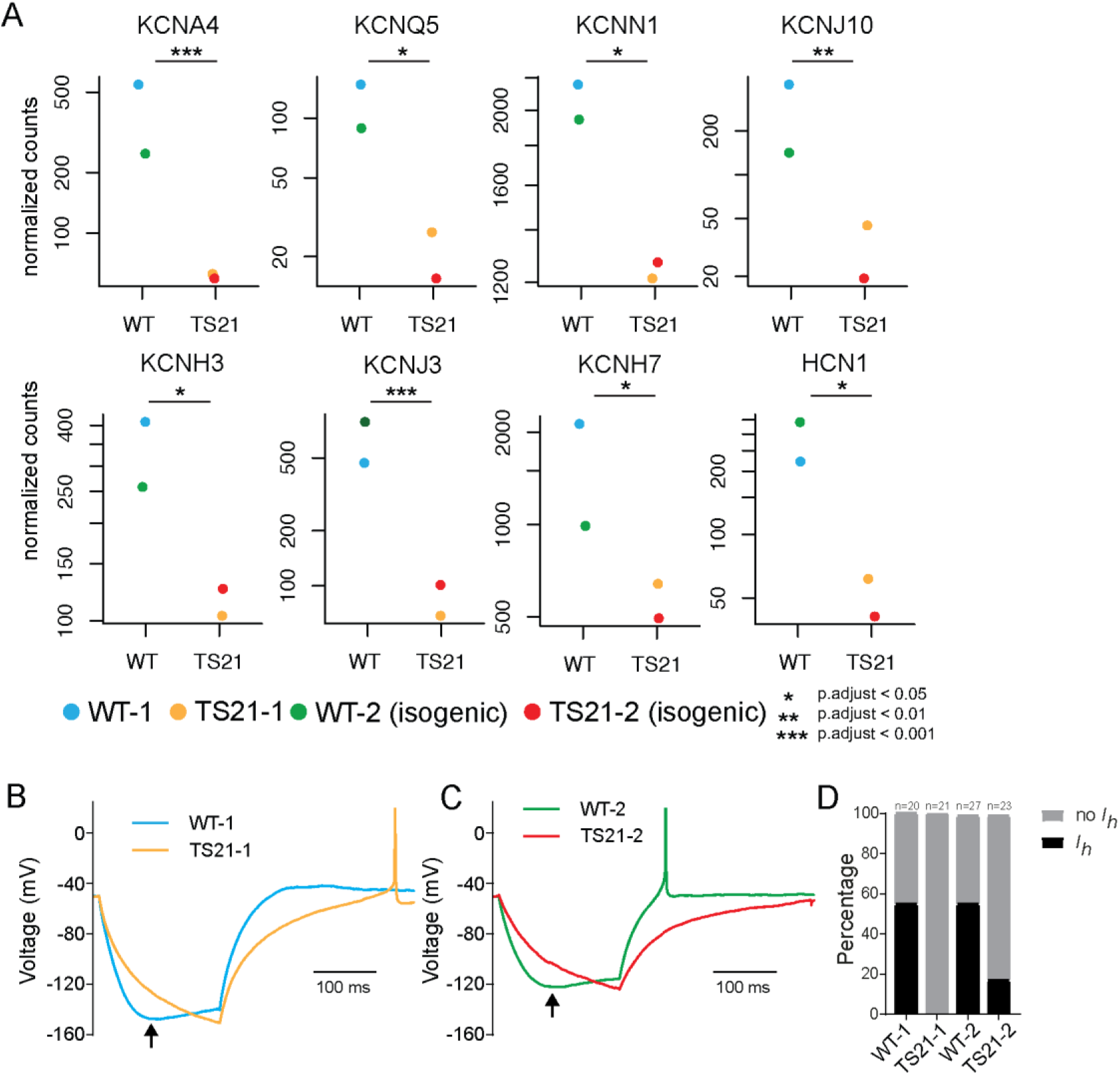
Decreased potassium channel expression in TS21 neurons. A: Gene expression analysis of K+ and K+/Na+ channel expression in TS21 and WT neurons, note that all 8 channels show a significant down-regulation in TS21 neurons (n = 1 sample from one neuronal induction per genotype). B, C: Detail of hyperpolarization activated currents elicited *in vivo* after current injection in current clamp mode. Arrows indicate the depolarising sag. D: Percentage of cells in which hyperpolarization activated currents (*I*_h_) could be measured per cell line. While 50% of WT neurons (26 out of 52) exhibited a clear sag potential, only 9.1% of TS21 neurons (4 out of 44) did so (WT-1 55.0% vs. TS21-1 0%, Fisher’s exact test ****P < 0.0001; WT-2 55.56% vs. TS21-2 17.39%, Fisher’s exact test ** P < 0.01).

We first asked if genes located on Hsa21 are among those significantly upregulated genes in TS21 neurons. Indeed, we found a significant upregulation of many Hsa21 located genes including Amyloid Precursor Protein (APP), Down Syndrome Cell Adhesion Molecule (DSCAM), the Glutamate Ionotropic Receptor Kainate Type Subunit 1 (GRIK1) and the Potassium Voltage-Gated Channel Subfamily J Member 6 (KCNJ6) (Figure 5—figure supplement 1A), which have all been implicated in DS pathology (Lana-Elola et al 2011, Wiseman et al 2009). Next, we asked if there are changes in the expression levels of the AMPA and NMDA receptor subunits. Whilst we did not detect changes in most AMPA and NMDA receptor subunits, we found a significant upregulation of GRIN2A in TS21 neurons. GRIN2A is located on Hsa16 and encodes for the NMDA type subunit 2A (NR2A), which is predominantly expressed in the adult brain, particularly in neurons (Law et al 2003, Sheng et al 1994). This increase in NR2A subunit expression is in line with a previous report that showed increased NR2A expression in a DS mouse model leading to changed Ca^2+^ dynamics in those neurons (Altafaj et al 2008) (Figure 5—figure supplement 1B, C).

Next, we performed a gene ontology (GO) analysis and found that the GO terms “synaptic transmission” and “Voltage gated potassium channel activity” were highly overrepresented amongst downregulated genes in TS21 neurons (Supplemental Table 1). This was reflected in a marked downregulation of voltage gated (KCNA4, KCNQ5, KCNJ10, KCNH3, KCNH7, KCNJ3) and calcium activated K+ channels (KCNN1) in TS21 neurons (Figure 5A). We also found a significant downregulation of the potassium/sodium hyperpolarization-activated cyclic nucleotide-gated channel 1 (HCN1), a mediator of rhythmic activity (Biel et al 2009). To further test the role of HCN1 in DS, we performed electrophysiological recordings of WT control and TS21 grafted neurons *ex vivo*. Intriguingly, we found that hyperpolarization-activated currents, elicited after current injection, were significantly reduced in TS21 neurons (Figure 5B-D). Our analysis further revealed a dramatic, near all-or-none loss of functional Ih current in the majority of TS21 neurons (Figure 5D). Those currents showed a strong correlation between membrane hyperpolarization and current voltage (Figure 5—figure supplement 3), suggesting that they are mediated by HCN channels, since membrane hyperpolarization is both necessary and sufficient to activate these channels. To further support the conclusion that this functional loss is driven by reduced HCN expression, we analyzed publicly available RNA-seq datasets from independent DS fetal brain cohorts and found a consistent downregulation of HCN1 (Figure 5—figure supplement 4A), corroborating our initial transcriptomic screen (Olmos-Serrano et al 2016). We also examined HCN expression in our own snRNA-seq atlas (Lattke et al 2026), in which we report widespread transcriptional dysregulation in excitatory neurons of the DS human fetal cortex. While these genes did not pass the stringent threshold for statistical significance (a known limitation of snRNA-seq for moderately expressed transcripts due to data sparsity), the data shows a clear trend of downward expression in DS excitatory neurons (Figure 5—figure supplement 4B), consistent with all our other findings.

## Discussion

In this study, we have longitudinally assayed the spontaneous neuronal network activity and connectivity of developing isogenic TS21 and control hiPSC-derived cortical forebrain neurons, both *in vitro* and *in vivo*. We found a modest decrease of spontaneous firing activity in TS21 neurons early in development, and a marked reduction (Martini et al 2021) of synchronized bursting activity in trisomic neuronal networks. Synchronized bursting is found in many different model systems (Corlew et al 2004, Khazipov et al 2004, Kilb et al 2011, Kirwan et al 2015, Leinekugel et al 2002, Robinson et al 1993, Winnubst et al 2015), suggesting a universal feature of neuronal network development and maturation. Interestingly, EEG recordings and event-related brain potentials of individuals with DS also exhibit altered cortical network activity and weaker cortical synchronization (Babiloni et al 2010, Karrer et al 1998, Velikova et al 2011), and decreased synchronization was also found in the neocortex (Cramer et al 2015) and in primary hippocampal neurons of DS mice (Stern et al 2015).

Our results suggest that the changes in synchronization observed in TS21 neurons are intrinsic to the excitatory neuronal network and unlikely to be mediated by the low number of interneurons (Bacci & Huguenard 2006) or astrocytes (Mizuno et al 2018), at least in these early stages of development. In support of this, we recently generated a single-cell multiomic atlas of the DS midgestational cortex that revealed broad changes in genomic programs, including downregulation of key neuronal differentiation genes involved in intellectual disability, primarily affecting the excitatory neuron lineage (Lattke et al 2026). These results have been recently validated by independent laboratories (Risgaard et al 2025, Vuong et al 2025) underscoring the significance of the mechanisms identified in the present study. Given the differences observed in the proportions of upper layer neurons and astrocytes generated from TS21 and control iPSCs in the xenotransplantation model 5 mpt (Figure 3—figure supplement 1), as also reported in human post-mortem brain (Klein & Haydar 2022), it is possible that cellular composition contributes to the differences in connectivity and/or network performance at these later developmental stages.

Our calcium imaging approach allowed us to quantify the activity of hundreds of individual neurons simultaneously and determine the proportion of active cells, an outcome that cannot be directly achieved with other methods such as Multi Electrode Arrays. While our current analysis of neuronal synchrony is based on mean pixel correlation, which provides a useful but coarse measure of network activity, future studies incorporating ROI-based approaches—such as cross-correlation or spike-time tiling coefficients—will be important to achieve a more precise, single-neuron level characterization of synchrony (Sintes et al, in preparation).

To investigate whether the reduction of synchronised neuronal network activity could result from altered connectivity between neurons, we used a retrograde, pseudotyped rabies virus (Brennand et al 2011, Kirwan et al 2015, Wickersham et al 2007), and found a significant decrease in connectivity between trisomic neurons. Importantly, trisomic neurons were not impaired in forming connections *per se*, nor was the virus spread inhibited by the extra Hsa21 copy. *In vivo,* rabies spread is tightly correlated with synaptic inputs and only occurs between chemical synapses and not via gap junctions or local spreading of the virus (Ugolini 2008), which raises the question of whether the lower connectivity rates in trisomic neurons could be explained by their reduced spontaneous activity. However, a complete block of spontaneous activity with TTX in control neurons did not alter the connectivity rate compared to untreated neurons, indicating that the decrease in connectivity observed in trisomic neurons is independent of neuronal activity. We also considered whether changes in synaptic numbers or functional properties could explain the deficit in connectivity in Ts21 neurons. A reduction of total synapse number has previously been reported in a DS mouse model (Chakrabarti et al 2007), but a study using hiPSC-derived cortical neurons did not detect differences in synapse numbers and further showed functional synapse formation in both trisomic and control neurons (Shi et al 2012b). Importantly, we also did not detect changes in the synaptic properties of trisomic neurons.

The finding of reduced connectivity between T21 cortical neurons is in line with a human study of MR functional connectivity showing a simplified brain network architecture in the DS brain, dominated by a high degree of local connectivity and a lack of long-range connectivity (Pujol et al 2015). Whilst we did not detect topological changes, we found a pronounced decline of interconnected neurons in trisomic networks, which could interfere with normal neuronal synchronization during development.

Our in vitro human neural models recapitulate a specific and critical window of early human corticogenesis, the period during which deep-layer excitatory neurons are generated. Notably, at the stages used for functional analysis (up to Day 60) our cultures do not contain significant numbers of SATB2+ neurons or GFAP+ astrocytes (Kirwan et al 2015, Shi et al 2012c). DS is a neurodevelopmental disorder with pathological origins in fetal life. Our model provides a window into the cellular and molecular deficits as they first emerge during this early developmental process, something that is impossible to study functionally in post-mortem tissue. Our findings align with and extend previous studies using alternative Down syndrome models, such as brain organoids (Wang et al 2025) and other hiPSC-derived systems. Organoid models have provided valuable insights into early neurodevelopmental phenotypes in DS, including altered interneuron proportions (Xu et al 2019), but also suggest that variability across isogenic lines can overshadow subtle trisomy 21 neurodevelopmental phenotypes (Czerminski et al 2022). However, these systems often lack the structural complexity, vascularization, and long-term maturation achievable *in vivo*. By using a xenotransplantation model, we were able to assess the maturation and functional properties of human neurons within a physiologically relevant environment over extended time frames, offering complementary insights into DS-associated circuit dysfunction (Huo et al 2018, Real et al 2018).

Finally, to further explore mechanisms of reduced synchronicity, we performed RNA sequencing of *in vitro* neurons. Our gene expression analysis found a strong upregulation of KCNJ6, as well as a downregulation of many voltage gated K+ channels, potentially implicating those channels in DS pathology. Potassium channels are important mediators of neuronal excitability and action potential shape. In particular, the K+ channel KCNJ6 is located on Hsa21 and leads to changes in synaptic plasticity in a DS mouse model (Cooper et al 2012). Furthermore, KCNJ6 currents are elevated in hippocampal neurons of DS mice due to impaired GABAergic inhibition (Best et al 2012). Single neurons from DS mice further exhibit reduced K+ currents related to changes in A-type, M-type and delayed rectifier channels (Stern et al 2015), which could impact network activity. We also found a downregulation of the HCN1 channel. HCN channels, which upon activation conduct a mixed Na^+^/K^+^ current, thus depolarizing the membrane towards firing threshold of action potentials, have been implicated in several neuronal functions, including the regulation of dendritic integration and synaptic transmission, as well as the generation of neuronal and network rhythmic oscillations in several brain regions (Biel et al 2009). Various HCN channel subtypes are differentially expressed at distinct developmental periods and contribute to network activity maturation (Bender & Baram 2008). In cortical neurons, hyperpolarization-activated currents are mainly mediated by the HCN1 subtype (Shah 2014) and HCN1 channels are important to regulate dendritic excitability (Tsay et al 2007), and misregulation can contribute to multiple neurological disorders (Chang et al 2019). Interestingly, Aβ production is dependent on HCN1 channel activity *in vitro* (Saito et al 2012) and a recent meta-analysis found a significant downregulation of HCN1 in the human AD brain (Haytural et al 2021), thus implicating HCN1 expression in AD pathology. Consistent with this evidence, we found a reduction of HCN-mediated hyperpolarization-activated currents in TS21 neurons *ex vivo*, which could potentially alter cortical network activity and development in TS21 neurons. However, further targeted functional validation will be needed to confirm a causal link.

In summary, our data from isogenic control and DS neurons reveal a fundamental difference in the development of DS excitatory neuronal networks and a direct influence of the extra Hsa21 copy in channel composition, neuronal network activity development and connectivity. Alterations in neural network synchronization and connectivity were observed in the excitatory neuron population, which may underlie some of the neurodevelopmental and neurological features of DS.

## Materials and Methods

### Generation of iPSCs

Skin biopsies were collected from donor individuals with DS by the University of Cambridge Dept. of Psychiatry, following approval of the Cambridgeshire Research Ethics Committee, with informed consent from the donor or representative. Samples were designated anonymous identifiers following collection. Skin punch biopsies were subdivided and cultured at 37°C in 5% CO2 in DMEM + Glutamax (ThermoFisher) supplemented with 10% fetal calf serum, penicillin (50 U ml^−1^), and streptomycin (50 μg ml^−1^) and 1mM Sodium Pyruvate (Sigma) until fibroblast outgrowth was observed. When plates were confluent, fibroblasts were trypsinized and transferred to T75 dishes for further expansion. Cellular reprogramming of fibroblasts was carried out using Cytotune 2.0 Sendai Reprogramming kits (ThermoFisher) according to the manufacturer’s protocol, and established using feeder free conditions in Essential-8 media (ThermoFisher).

### Cortical neuron differentiation

All cells were maintained at 5% CO2 at 37°C in a humidified incubator. WT1: NDC1.2(Israel et al 2012), DS1: DSIPS(Park et al 2008), WT2: (Tho3F), DS 2: (Tho3B) (Real et al 2018) iPSC were cultured in Essential-8 medium (ThermoFisher) as feeder free cultures on Geltrex (ThermoFisher) coated plates. Neuronal induction was performed as described previously(Shi et al 2012a). In brief, iPSCs were harvested with Accutase (Innovative Cell Technologies) as single cell suspension and plated at a density to reach 100% confluency in 24h and neuronal induction was started (day 0). Essential-8 medium was changed to neuronal induction medium consisting of neuronal maintenance medium supplemented with 10μm SB43152 (Tocris) and 1μm Dorsomorphin (Tocris). The medium was changed daily until day 12. On day 12 the neuroepithelial sheet was lifted with Dispase (ThermoFisher), broken into smaller clumps and plated on laminin (Sigma) coated plates in neuronal induction medium. The 1mg ml^-1^ laminin stock solution was diluted 1:100 with PBS to a final concentration of 10 μg ml^-1^. Plates were covered with laminin and incubated for 4h at 37°C. Laminin solution was aspirated before addition of the medium. On day 13 the medium was changed to neuronal maintenance medium supplemented with 20 ng ml^-1^ FGF2 (PeproTech). Medium was changed every other day and FGF2 was withdrawn from the medium on day 17. Cells were split with Dispase at a 1:2 ratio when neuronal rosettes started to meet. On day 25 cells were dissociated with Accutase (ThermoFisher) and re-plated on Poly-L-ornithine (Sigma Aldrich) / laminin coated plates. The plates were covered with 0.01% (wt/vol) poly-L-ornithine solution and incubated for 4h at 37°C. Plates were washed twice and coated with laminin as described above. Until day 34 cells were expanded at a 1:2 ratio when they reached 90% confluency. On day 36 neurons were plated on laminin coated plates at 85000 cells/cm^2^ and used for subsequent experiments.

Neuronal maintenance medium (NMM) (1L) consists of 500ml DMEM:F12+glutamax (ThermoFisher), 0.25ml Insulin (10mg ml^-1^, Sigma), 1ml 2-mercaptoethanol (50mM ThermoFisher), 5ml Non-essential amino acids (100X ThermoFischer), 5ml Sodium Pyruvate (100mM, Sigma), 2.5ml Pens/Strep (10000U/μl, ThermoFisher), 5ml N2 (ThermoFisher), 10ml B27 (ThermoFisher), 5ml Glutamax (100X, ThermoFisher) and 500ml Neurobasal (ThermoFisher) medium.

### Immunofluorescence staining

Cells were washed with PBS and fixed with 4% paraformaldehyde for 10min at room temperature (RT). They were washed (5min 3 X TBS, 3 X TBS + 0.3% Triton-X100 at RT) and blocking was performed with 5% normal goat serum in TBS + 0.3% Triton-X100 for 1h at RT. Cells were incubated with primary antibodies overnight at 4C using the following antibodies: TBR1 (1:1500, Abcam: ab31940), CTIP2 (1:1000, Abcam: ab18465), TUJ1 (1:1000, Biolegend: 8012021), GFP (1:1000, Abcam ab13970), mCherry (1:1000, Clontech: 632543). On the next day, cells were washed (5min 3 X TBS, 3 X TBS + 0.3% Triton-X100 at RT) and incubated for 1h at RT with Alexa-dye conjugated secondary antibodies (Thermo Scientific) diluted 1:500 in TBS + 0.3% Triton-X100. Cells were washed 6 X for 5min with TBS and 4’,6-Diamidino-2-Phenylindole (DAPI, 5μg ml^-1^ final concentration) was added to the second wash step. Imaging was performed using an inverted Leica TCS SP8 scanning confocal microscope (Leica Microsystems) or Opera Phenix High-Content Screening System (Perkin Elmer).

### *In vitro* calcium imaging

For longitudinal calcium activity time series one (WT-2) or two (WT-1, TS21-1, TS21-1) independent neuronal differentiations were analysed. Neurons were switched to complete BrainPhys (BP) medium (STEMCELL Technologies) at day 40 and network activity was assessed between day 50 and day 80. 6 to 8 independent wells were assayed per line and the same wells were assayed over this time period. Neurons were loaded with the calcium indicator Oregon Green 488 BAPTA-1, AM (OGB, ThermoFisher). OGB was dissolved in DMSO and 0.4% w/v Pluronic F-127 (Sigma-Aldrich) to a concentration of 0.8mM OGB. This solution was further diluted with BP medium to a final concentration of 3.2μm and neurons were incubated with OGB for 1h at 37°C and 5%CO_2._ Neurons were washed 3 times with BP medium and network activity was assayed 30min afterwards. The cells were imaged at 37°C and 5%CO_2_ on a Deltavision wide field fluorescence microscope (GE Healthcare Lifescience) equipped with an EMCCD camera. Four two-minute time lapse movies at 10Hz were captured per well. Time lapse movies were analyzed using a customwritten Matlab (Mathworks) script (Catran2). The average number of analysed cells per ROI were WT-1 142±15, WT-2 140±24, TS21-1 109±41, TS21-2 136±16 mean±SD. For pharmacological experiments cells were recorded in BP medium containing 1μM tetrodotoxin (TTX, Tocris Bioscience), 50μm 6-cyano-7-nitroquinoxaline-2,3-dione (CNQX, Tocris Bioscience), or 50μm DL-2-Amino-5-phosphonopentanoic acid (AP5, Tocris Biocience). After the recording session drugs were removed by three washes with BP medium and the same locations were imaged 45min later.

### *In vitro* Calcium Imaging Time-Lapse Analysis

Regions of interest (ROIs) were manually drawn in ImageJ on the mean z-projected image from each calcium imaging video. Separate ROIs were defined for neuronal somata and background areas to allow baseline correction, ensuring that fluorescence signals reflected true cellular activity rather than background noise. Raw movies were motion-corrected using the CaImAn Python package (Giovannucci et al 2019). For each ROI, raw fluorescence (F) was extracted, and normalized signals were obtained by subtracting mean background fluorescence, yielding a background-corrected ΔF trace. Normalized fluorescence data for each ROI were exported to Excel for downstream analysis.

- **Maximum Amplitude** For each video, ΔF traces from all cell ROIs were averaged to generate a population activity trace. The maximum amplitude of this trace was extracted per video, providing one representative amplitude value for group comparison (WT vs. TS21).
- **Burst Detection** Bursts were defined as transients exceeding a cell-specific threshold of μ+3σ, where μ is the mean baseline fluorescence and σ the baseline standard deviation. This stringent criterion minimized false positives from noise. To separate independent events, a minimum inter-burst interval (IBI) of 0.5 s was applied, consistent with the temporal resolution of OGB-1 (∼500 ms).
- **Burst Frequency** Burst frequency was quantified as average across all ROIs, capturing overall network activity; For each ROI, we quantified burst number, onset times, amplitudes, and frequencies, with results systematically saved in Excel for further analysis. All stage specific analyses were performed at day 60 of differentiation, the stage previously identified as showing the greatest divergence in synchronised activity between WT and TS21 cells.
- **Statistical Analysis** Group differences in burst amplitude and frequency were assessed using the Mann–Whitney U test. Effect sizes were reported as Cohen’s *d*. Statistical significance was defined as *p* < 0.05, while *p* ≥ 0.05 was considered not significant (*ns*).

### *In vitro* electrophysiological recordings

Cortical neurons were incubated with artificial cerebral spinal fluid containing 125 mM NaCl, 25 mM NaHCO3, 1.25 mM NaH2PO4, 3 mM KCl, 2 mM CaCl2, 25 mM glucose, and 3 mM pyruvic acid and bubbled with 95% O2 and 5% CO2. Borosilicate glass electrodes with resistance of 6–10 MΩ were filled with an artificial intracellular solution containing 135 mM potassium gluconate, 7 mM NaCl, 10 mM HEPES, 2 mM Na2ATP, 0.3 mM Na2GTP, and 2 mM MgCl2, and positioned over a cortical neuron to form a whole-cell patch. Recordings were made using a Multiclamp 700A amplifier (Molecular Devices), and signals were sampled and filtered at 20 kHz and 6 kHz, respectively. A low-pass Gaussian filter was applied to filter out high-frequency noise.

### Lentiviral generation

Third generation lentivirus was produced by calcium phosphate transfection of the Lenti-X 293T cell line (Clontech) with pBOB-synP-HT (Addgene plasmid # 30456) expressing a histone:GFP, TVA and rabies glycoprotein separated by 2A sites from the human synapsin promoter. Cells were co-transfected with the packaging plasmids plasmids pRSV-Rev (Addgene plasmid # 12253) and pMDLg/pRRE (Addegene plasmid # 12251) and the VSVG envelope plasmid pMD2.G (Addgene plasmid # 12259). Medium was replaced 16h after transfection. The virus supernatant was harvested after 8h replaced with new medium and harvested again after 12h. Both collections were pooled, debris was removed with a 0.45μm filter and virus supernatant was stored at –80°C for subsequent use.

### Monosynaptic rabies tracing

Neurons were grown in BP medium as described above. On day 40 they were infected with low titer SYN-HTG lentivirus. Medium was exchanged the next day to remove the lentivirus. On day 50 neurons were infected with mCherry expressing pseudotyped rabies virus provided by Marco Tripodi, MRC Laboratory of Molecular Biology, Cambridge UK. Neurons were fixed 6 days later and immunofluorescence stainings for GFP and mCherry were performed as described above. To test the effects of neuronal activity on virus spread, neurons were infected with low titer SYN-HTG lentivirus on day 38. Rabies infection was performed on day 48. 24h after the infection 1μM TTX was added to the medium. Medium was changed every other day with TTX supplemented until fixation on day 54.

### RNAseq library preparation and analysis

Neurons were grown in neuronal maintenance medium. On day 50 they were harvested with Accutase, homogenized with a QIAshredder (Qiagen) and RNA was isolated using the RNeasy Mini Kit (Qiagen) following the manufacturer’s protocol. RNA quality was assayed on a 2200 TapeStation Instrument (Agilent) and sequencing libraries were prepared using the TruSeq Stranded Total RNA Library Prep Kit (Illumina) following the manufacturer’s protocol. 50bp single read ends were performed on a HiSeq4000 (Illumina) sequencer. Each sample library comprised 91-109 million reads, of which ∼83% were mapped to the GRCh37 genome using STAR (Trapnell et al 2009). DESEq2 (Love et al 2014) was run on counts data for genes with >1 count. Differentially expressed (DE) genes were selected using a Benjamini-Hochberg (BH) adjusted p-value of 0.05; for Figure 5—figure supplement 2, the gene counts are normalized for library size using DESeq2. Goseq (Young et al 2010) was used for gene ontology (GO) analysis of DE genes. The list of downregulated or upregulated genes were compared individually against the list of all detected genes and significantly overrepresented GO terms were selected using a BH adjusted p-value of 0.05.

### Human iPSC-derived Neuron Transplantation

Animal experiments were conducted in accordance with the Animals (Scientific Procedures) Act 1986, UK. Immunodeficient mice (JAX™ NSG, Charles River) aged between 3-5 months were given 3% isoflurane mixed with oxygen as induction anaesthesia, followed by an intraperitoneal injection of ketamine (0.083 mg/g) and xylazine (0.0078 mg/g). Craniotomies were performed over the right somatosensory cortex, as previously described (Holtmaat et al 2005). Neural cortical progenitors were allowed to differentiate for 31 days, before being transduced with a lentiviral vector carrying either GFP or GCaMP6s plus tdTomato genes, under the human Synapsin-1 promoter. Between days 35 and 38, cells were dissociated and injected into the right somatosensory cortex with a glass needle using a micro syringe pump (UMP-3, World Precision Instruments), at the following stereotactic coordinates: anterior-posterior = –1.8 mm; medial-lateral = +2.8 mm; dorsal-ventral = –0.5 mm from bregma. Approximately 40.000 cells were injected in 1 µl of sterile cortex buffer solution. A 5-mm diameter glass coverslip was placed over the craniotomy and sealed with cyanoacrylate tissue adhesive. The exposed skull was covered in dental cement and a metal plate placed on the left side of the skull, for positioning at the 2-photon microscope stage.

### *In vivo* calcium Imaging

Animals (same cohort as in (Real et al 2018)) were imaged in a custom built 2-photon microscope (Prairie Technologies) equipped with a tunable Ti:saphire laser (Coherent) and a 20x water immersion objective (NA 1.2 Olympus). Animals were tightly head-fixed to the microscope stage and lightly anesthetized using isoflurane at a concentration between 0.5-1% mixed with oxygen. Body temperature was maintained by placing the animals on a heat pad. Transplanted cells were first identified by their tdTomato signal using a 1040 nm laser beam, and single plane images were taken. GCaMP6 was then excited using a 920 nm pulsed laser beam, and five-minute time lapse movies were acquired at 3 Hz in a single plane. Data was acquired for approximately 60 minutes per animal. Time lapse movies were corrected for possible within-scan motion using the moco plugin in ImageJ (NIH). A pixel-wise correlation analysis was performed on all calcium movies in the following way: a median projection over time of the calcium imaging movie was subtracted from each frame in the movie to provide a ΔF time-series. The ΔF time-series of each pixel was then correlated with every other pixel in the movie. The mean and the standard-deviation of the distribution of correlation values was then taken for each pixel.

### *Ex vivo* Electrophysiology

Acute coronal brain slices (350 µm) were maintained at room temperature (22-25 °C) in external solution (NaCl 125 mM, KCl 2.5 mM, NaH_2_PO_4_ 1.25 mM, NaHCO_3_ 25 mM, CaCl_2_ 2 mM, MgCl_2_ 1 mM, D-glucose 10 mM and D-mannitol 15 mM, pH adjusted to 7.4 and equilibrated with 95% O_2_ and 5% CO_2_). Human neurons were identified in the somatosensory cortex by excitation of GFP. Whole-cell current-clamp recordings were performed at 35 °C using borosilicate glass micropipettes with a resistance of 4-8 MΩ filled with intracellular solution (K^+^ gluconate 130 mM, KCl 10 mM, EGTA 0.5 mM, NaCl 1 mM, CaCl_2_, 0.28 mM, MgCl_2_ 3 mM, Na_2_ATP 3 mM, GTP 0.3 mM, phosphocreatine 14 mM and HEPES 10 mM, pH adjusted to 7.2). Current-voltage relationships were evoked by stepwise current injection from –80 up to +50 pA in steps of 10 pA.

### Immunohistochemistry

At the end of the *in vivo* experiments, animals were given an overdose of ketamine and xylazine, and were intracardially perfused with 4% paraformaldehyde (PFA). Brains were dissected and fixed overnight in 4% PFA solution. 40 µm-coronal brain sections were cut using a vibratome (Leica Biosystems) and stained free-floating. Briefly, sections were permeabilized and blocked in a 10% donkey serum in PBS with 0.5% Triton X-100 for 1h at room temperature, followed by overnight incubation at 4 °C with anti-human nuclei antibody (1:200, Millipore: MAB 1281), anti-Tbr1 antibody (1:200, Abcam: ab31940), SATB1/2 (1:200, Abcam: ab51502), Pax6 (1:300, Biolegend: 901301), Ki67 (1:1000, Abcam: ab15580), GFAP (1:1000, DAKO: Z0334) in 5% donkey serum in PBS with 0.1% Triton X-100. Sections were subsequently washed and incubated for 2h at room temperature with Alexa Fluor conjugated secondary antibodies (1:1000, Thermo Fisher). Nuclei were counterstained using 4’,6-diamidino-2-phenylindole (DAPI, Thermo Fisher Scientific), before sections were washed and mounted onto glass slides using ProLong® Gold Antifade Mountant (Thermo Fisher Scientific). Sections were imaged with an inverted scanning confocal TCS SP8 microscope (Leica Microsystems) using a 40x objective (NA 1.45).

## Acknowledgments

The authors thank individual donors for providing skin biopsies; Dr Vickie Stubbs and Ellie Tuck of the Livesey lab for technical advice, and Kay Harnish of the Gurdon Institute for preparation of cDNA libraries; E. Mustafa, A. Czerniak, K. Horan, A. Matthews, and E. Rowley for assistance with animal care and monitoring. We thank Marco Tripodi (MRC, Cambridge UK) for the kind gift of the pseudotyped rabies virus and members of the Livesey and De Paola labs for feedback and support. We thank Michael Lattke (Imperial College London) for providing the HCN expression data of Figure 5, figure supplement 4).

This work was supported by a Wellcome Senior Investigator award (F.J.L.); the Alborada Trust of the ARUK Stem Cell Research Centre (M.P. and F.J.L.); Great Ormond Street Children’s Charity (Stem Cell Professorship, F.J.L.). the UK Medical Research Council and Ministry of Health, Singapore (V.D.P.); the GABBA Ph.D. program (FCT fellowship PD/BD/52198/2013), the Rosetrees Trust, and Alzheimer’s Research UK (R.R.); the UK Dementia Research Institute (grant code DRIImp17/18 Q3 to S.J.B.); and the Medical Research Council (grant code MRC.A654.5QB40, to M.A.S.).

## Author contribution

F.J.L and M.P. conceived, planned and analyzed the *in vitro* aspects of the manuscript. V.D.P and R.R. conceived and planned the live imaging, characterization, and analysis of transplanted donor-derived neurons and contributed to the study design; M.P performed and analysed iPSC differentiations, immunohistochemistry, RNA extractions, calcium imaging experiments, pseudotyped rabies virus labelling, *in vitro* electrophysiology recordings, and prepared the relevant figures. A.S. analyzed the gene expression and prepared the relevant figures. H.R. developed and provided calcium imaging analysis scripts.

R.R. and V.D.P. performed the grafting and the two-photon imaging experiments, analyzed the data, and prepared the related figures and text; M.A.S. performed *ex vivo* electrophysiology recordings and analysis from graft tissue generated by V.D.P. and R.R.; T.D. analysed the *in vitro* calcium imaging data with input from V.D.P.; S.J.B. analyzed the *in vivo* calcium imaging data and prepared the relevant figures and text with input from R.R. and V.D.P.; F.J.L led the project; and F.J.L and M.P. wrote the paper with contributions from all authors. All authors approved the final manuscript.

## Figures

**Figure 1—figure supplement 1.**
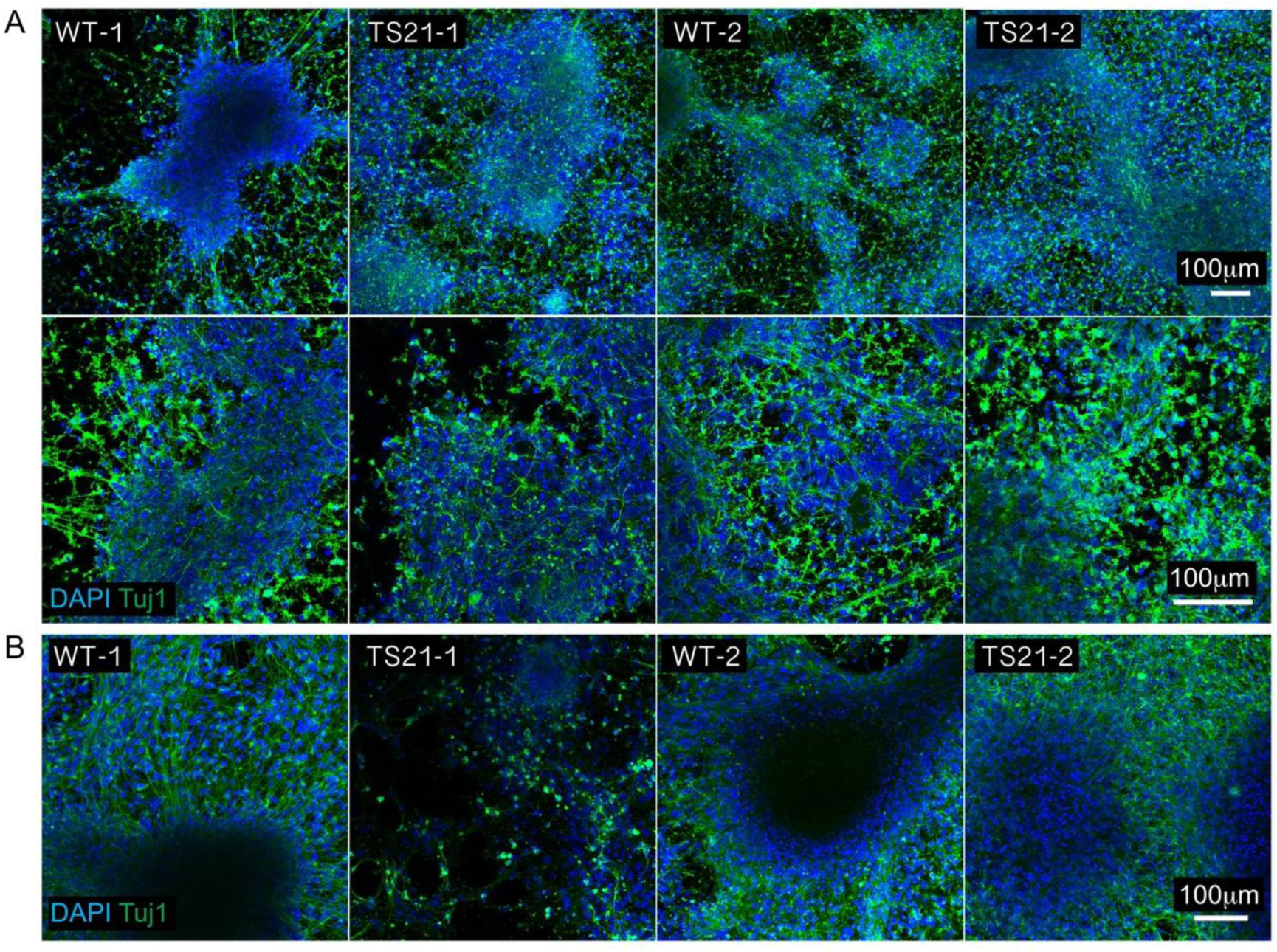
Confocal images of neuronal cultures from two control (WT-1, WT-2) and two trisomic (TS21-1, TS21-2) donors. A: Confocal image stacks of WT and TS21 trisomic neurons at day 50. Immunofluorescence staining for the pan neuronal marker Tuj1 and DAPI. B: Confocal image stacks of WT and TS21 neurons at day 70. Immunofluorescence staining for Tuj1 and DAPI.

**Figure 1—figure supplement 2.**
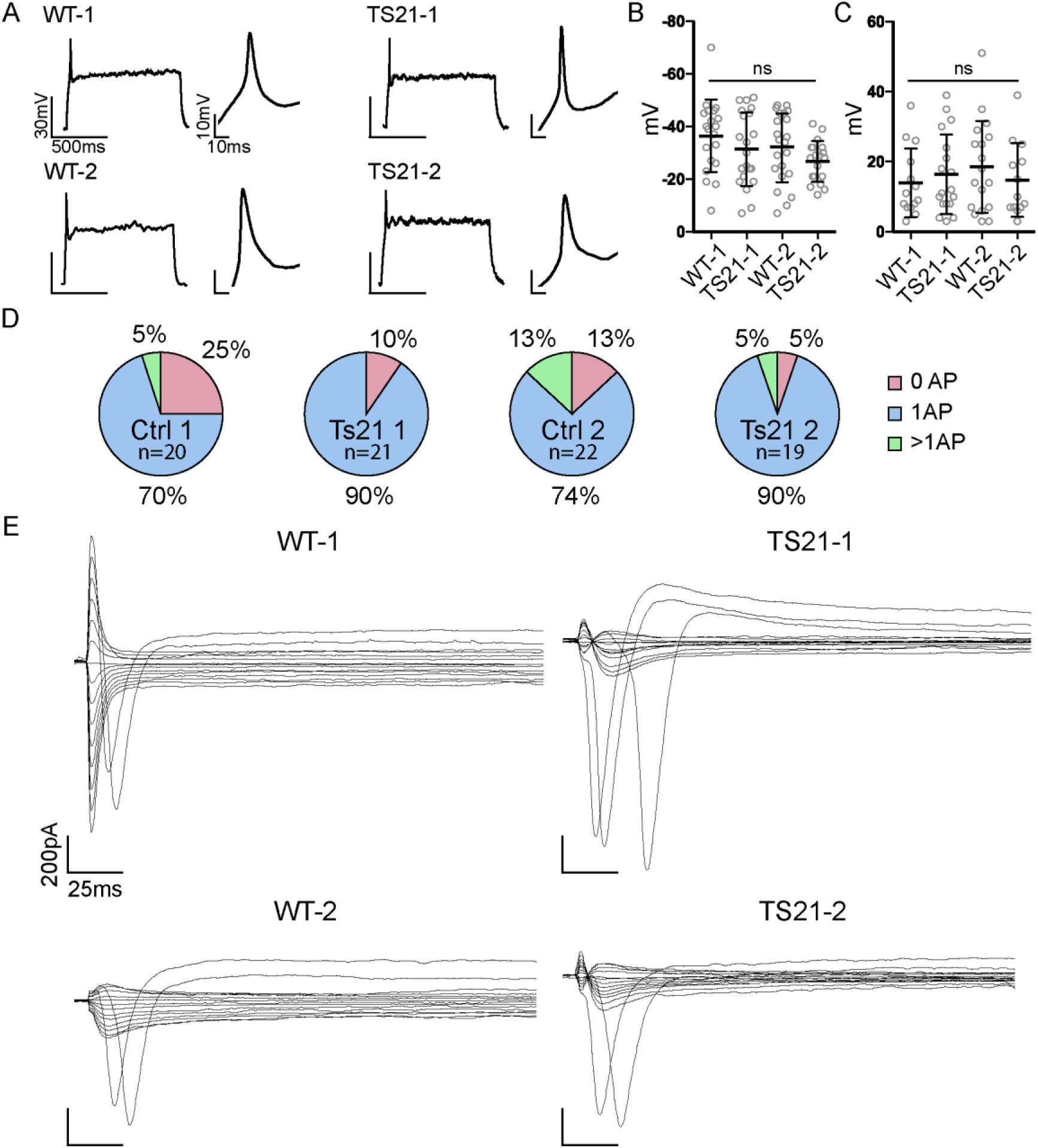
Electrophysiological characterization of TS21 and WT control neurons at day 53/54 shows unaltered whole cells properties in trisomic neurons. A: Whole cell electrophysiological recordings of TS21 and WT neurons at day 53/54. Neurons fire a single action potential (AP) after current injection. Inset shows a higher magnification of the AP. B: Resting membrane potential (WT-1: n=20, –36 ±14mV; TS21-1: n=21, –31 ± 14mV; WT-2: n=22, –32 ± 13mV; TS21-2: n=19, –27 ± 8mV; mean±SD, one-way ANOVA, data is from one neuronal induction per genotype), C: AP amplitude after current injection (WT-1: n=14, 14 ± 10mV; TS21-1: n=18, 16 ± 11mV; WT-2: n=17, 19 ± 13mV; TS21-2: n=13, 15 ± 11mV; mean±SD, one-way ANOVA, data is from one neuronal induction per genotype). D: Number of action potentials fired after current injection in current clamp mode. E: normal Na+ currents in TS21 and control neurons when voltage clamped.

**Figure 2—figure supplement 1.**
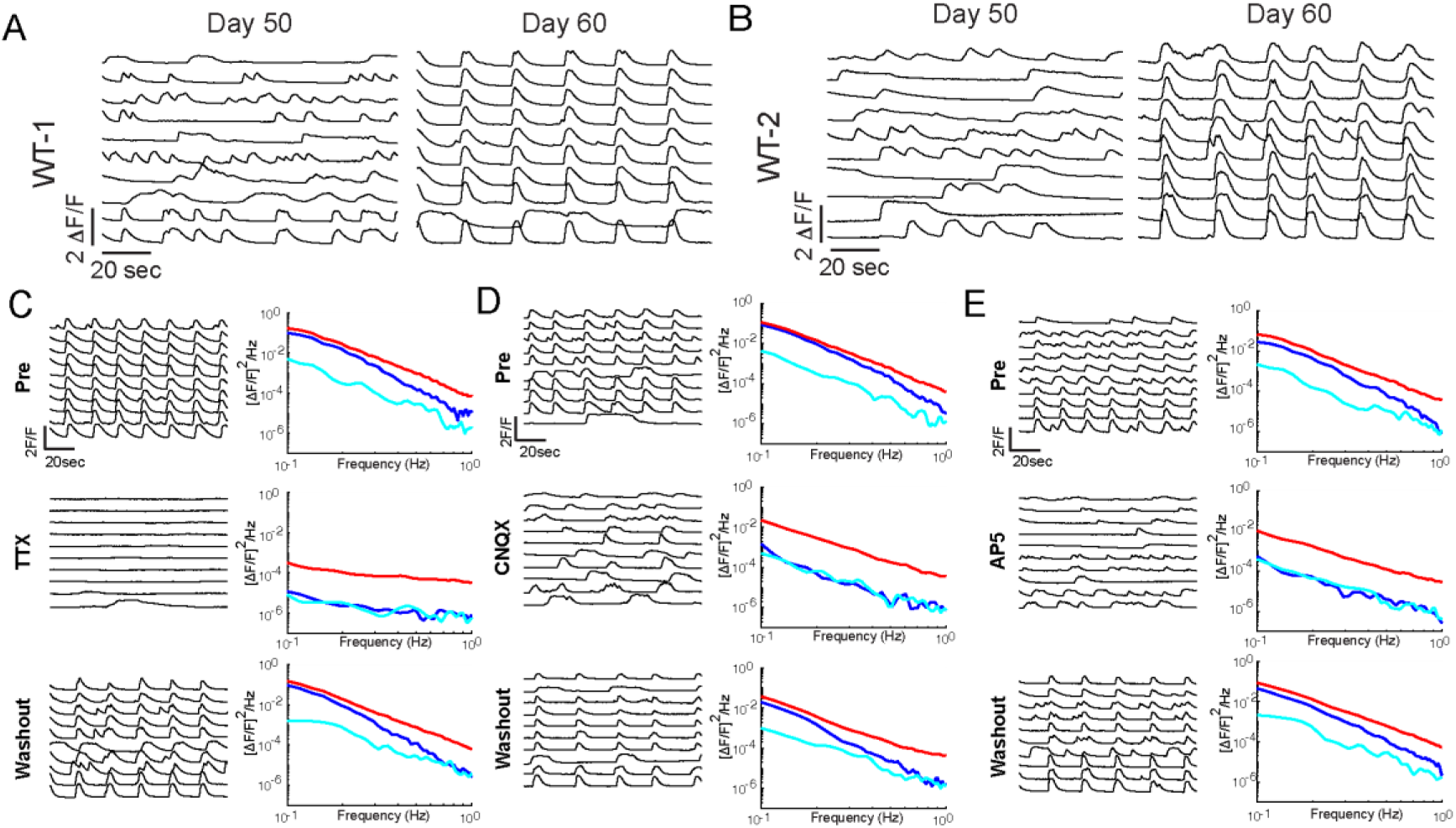
Spontaneous synchronized network activity development is AMPA and NMDA receptor dependent. Traces of fluorescence changes (ΔF/F) in WT neurons loaded with the calcium indicator OGB at day 50 and 60. A: WT-1, B: WT-2 (disomic isogenic). C, D, E: Left: Synchronized network activity is NMDA and AMPA receptor dependent. WT-1 neuronal networks were treated with three different drugs, during their synchronized phase. Pre: activity before pharmacological treatment, drugs (TTX, AP5, CNQX), Washout: 1h after pharmacological treatment. Left: ΔF/F traces of single neurons loaded with OGB, Right: corresponding power spectra of the 50 most active neurons, red line: average power of all neurons, blue line: power spectrum of the network, cyan line: randomized network power. C: TTX, synchronized network activity is reversely blocked by the Na channel blocker TTX, D: CNQX, synchronized network activity is reversely blocked by the AMPA receptor antagonist CNQX. E: AP5, synchronized network activity is reversely blocked by the NMDA receptor antagonist AP5.

**Figure 2—figure supplement 2.**
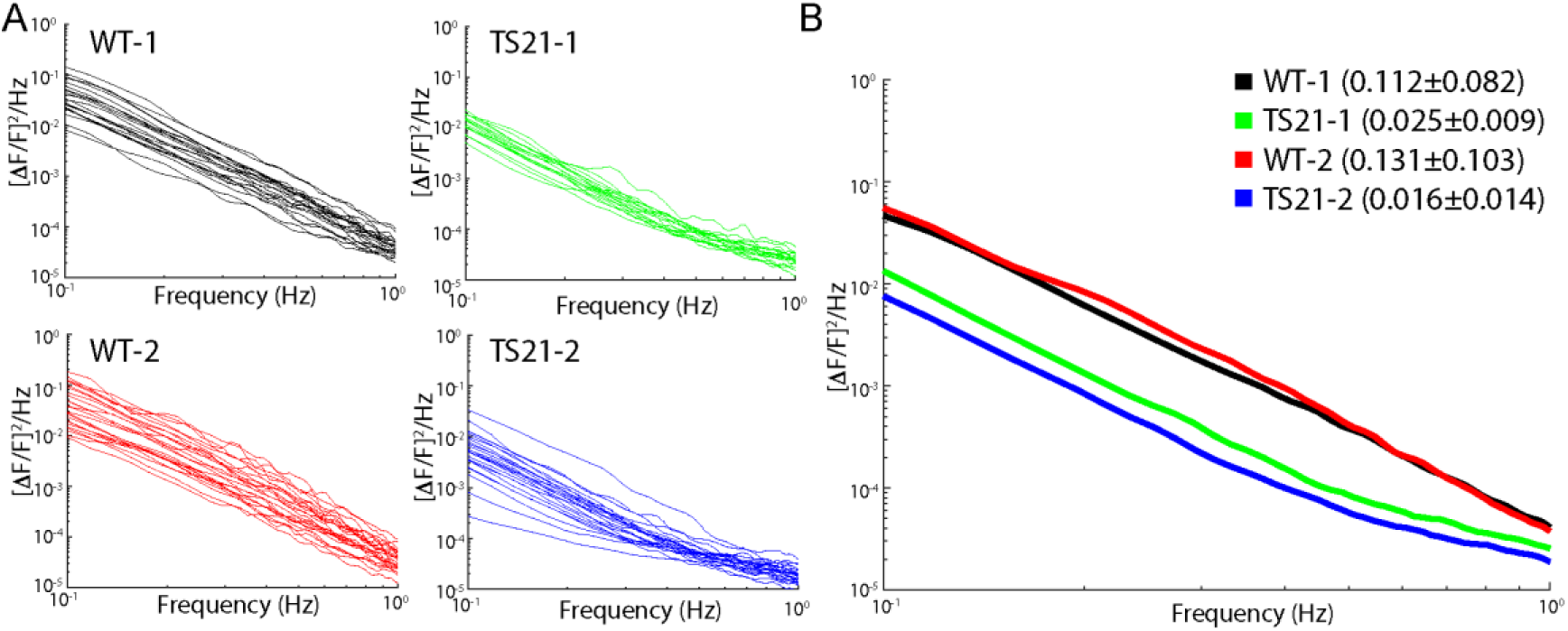
Reduced activity in trisomic TS21 neuronal networks at day 50. A: Average power spectra of TS21 and WT control neurons. B: averaged neuronal power spectra of traces shown in A. Note the decreased activity in both trisomic TS21 lines compared to WT, control neurons. Values show the mean integral of the power spectra over 0.1 and 1 Hz (WT-1 vs. WT-2: ns, WT-1 vs. TS21-1: **, WT-1 vs. TS21-2: ***, WT-2 vs. TS21-1: ***, WT-2 vs. TS21-2: ***, TS21-1 vs. TS21-2, ns, Tukey’s multiple comparisons test, *P <0.05), WT-1 n = 22 ROIs, TS21-1 n = 17 ROIs, WT-2 n = 25 ROIs, TS21-2 n = 24 ROIs).

**Figure 2—figure supplement 3.**
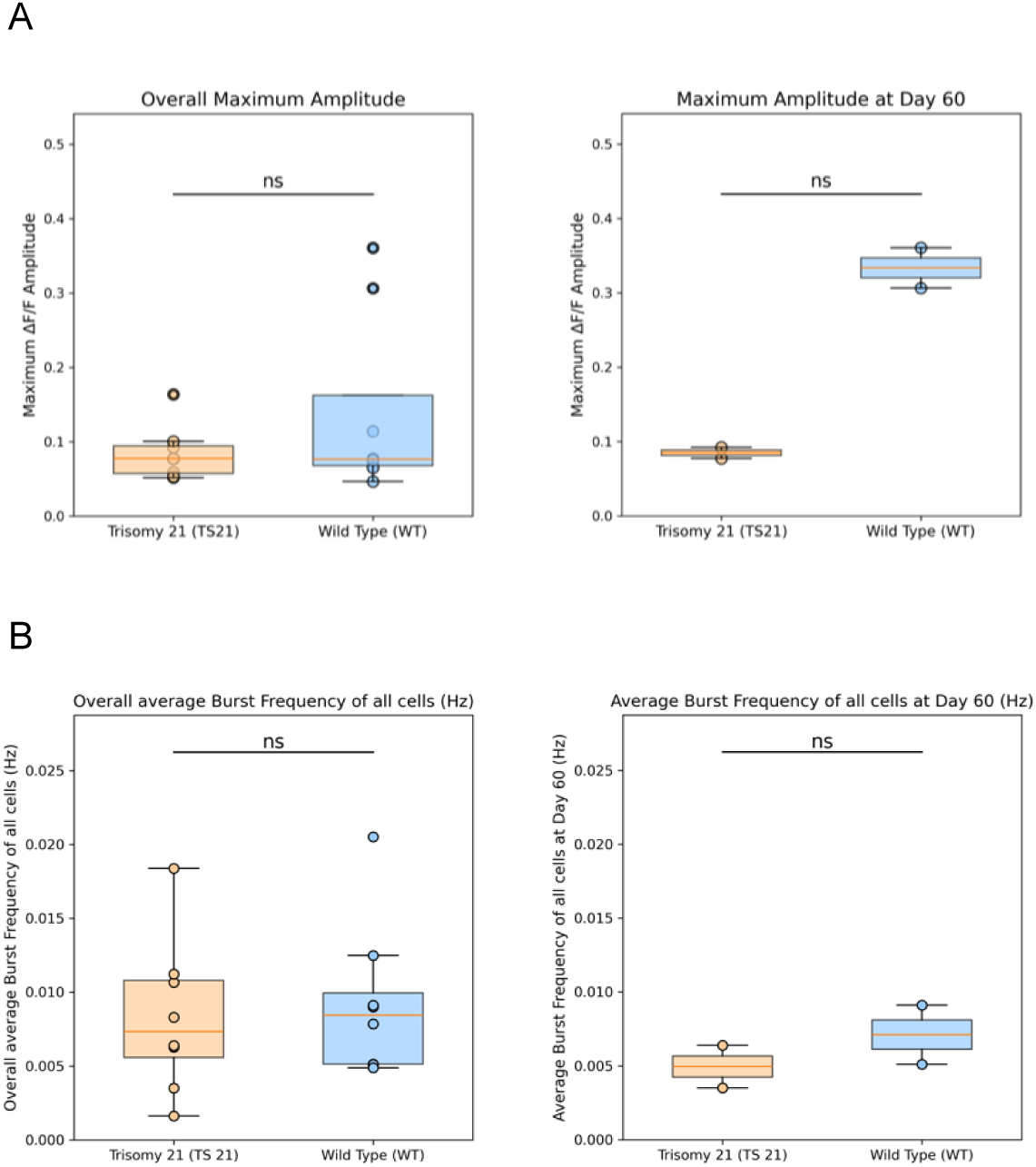
Maximum amplitude and burst frequency from *in vitro* calcium imaging in WT and Ts21 cells. (A) Maximum amplitude of calcium transients averaged across neuronal populations in WT and Ts21 cultures (Left, n = 8 WT, 8 TS21; Mann Whitney U_p = 0.72, Cohen’s d = –0.61; right, n = 2 WT, 2 TS21, Mann Whitney U_p = 0.33, Cohen’s d = –8.83). (B) Burst frequency quantified across ROIs in WT and Ts21 cultures (Left, n = 8 WT, 8 TS21; Mann Whitney U_p = 0.80, Cohen’s d = –0.19; right, n = 2 WT, 2 TS21, Mann Whitney U_p = 0.67; Cohen’s d = –0.88).

**Figure 3—figure supplement 1.**
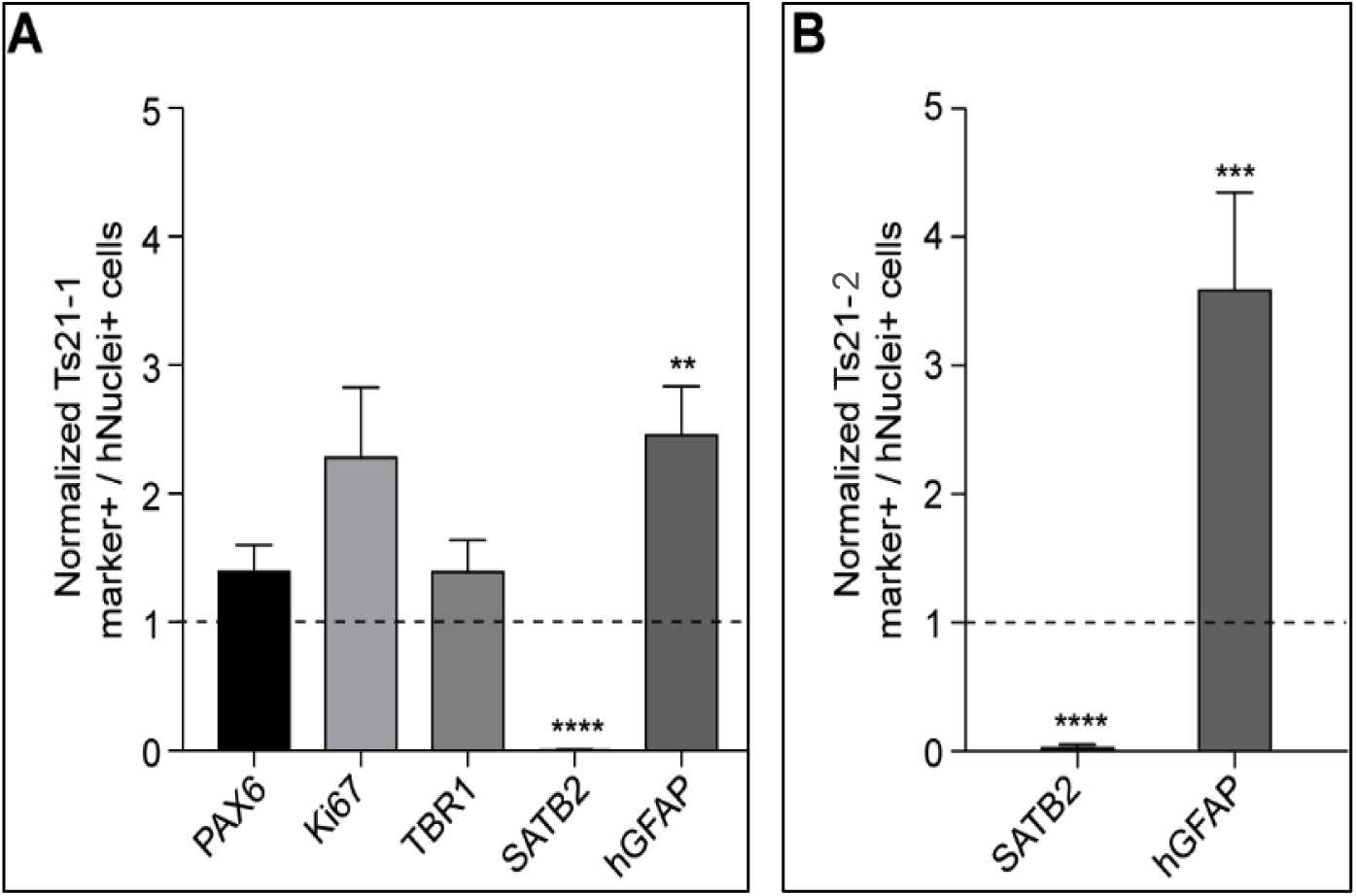
SATB2 expression is significantly reduced in TS21 grafts at 5 mpt. (A) Quantification of marker+ / hNuclei+ cells in Ts21-1 grafts (n = 3 mice, mean number of ROIs sampled per cell marker = 67) normalized to WT-1 (dashed line, n = 3 mice, mean number of ROIs per cell marker = 37). (B) Quantification of SATB2+ / human nuclei+ cells and GFAP+ / human nuclei+ cells in TS21-2 grafts (n = 2 mice, mean number of ROIs per cell marker = 22) normalized to WT-2 (dashed line, n = 4 mice, mean number of ROIs per cell marker = 45). Mann-Whitney U-test, **P < 0.01, ***P < 0.001, ****P < 0.0001. Data are shown as mean ± SEM.

**Figure 3—figure supplement 2.**
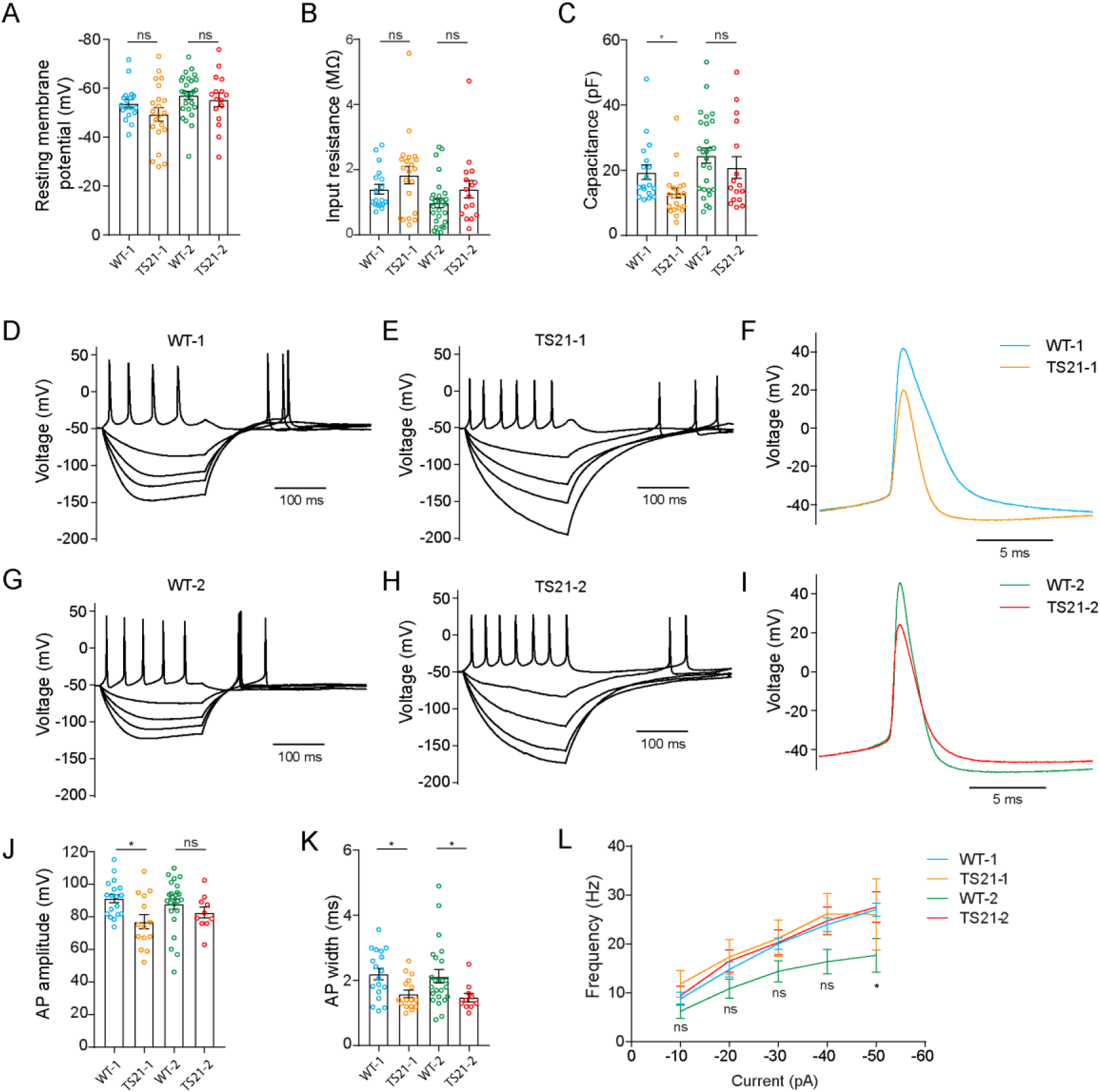
*Ex vivo* electrophysiological characterisation of TS21 and WT neurons. A: Resting membrane potential. Unpaired two-tailed t-test; ns, not significant. B: Input resistance. Mann-Whitney U-test; ns, not significant. C: Capacitance. Mann-Whitney U-test, **P <0.01; ns, not significant. (D-E, G-H) Representative current clamp recordings of membrane potentials after current injection in WT-1 (D), TS21-1 (E), WT-2 (G) and TS21-2 (H) neurons. F: Detail of an action potential in WT-1 and TS21-1 neurons. I: Detail of an action potential in WT-2 and TS21-2 neurons. J: Action potential amplitude. Mann-Whitney U-test, **P <0.01; ns, not significant. K: Action potential width. Mann-Whitney U-test, *P <0.05. L: Frequency of action potentials in response to positive current injection. Error bars represent SEM.

**Figure 4—figure supplement 1.**
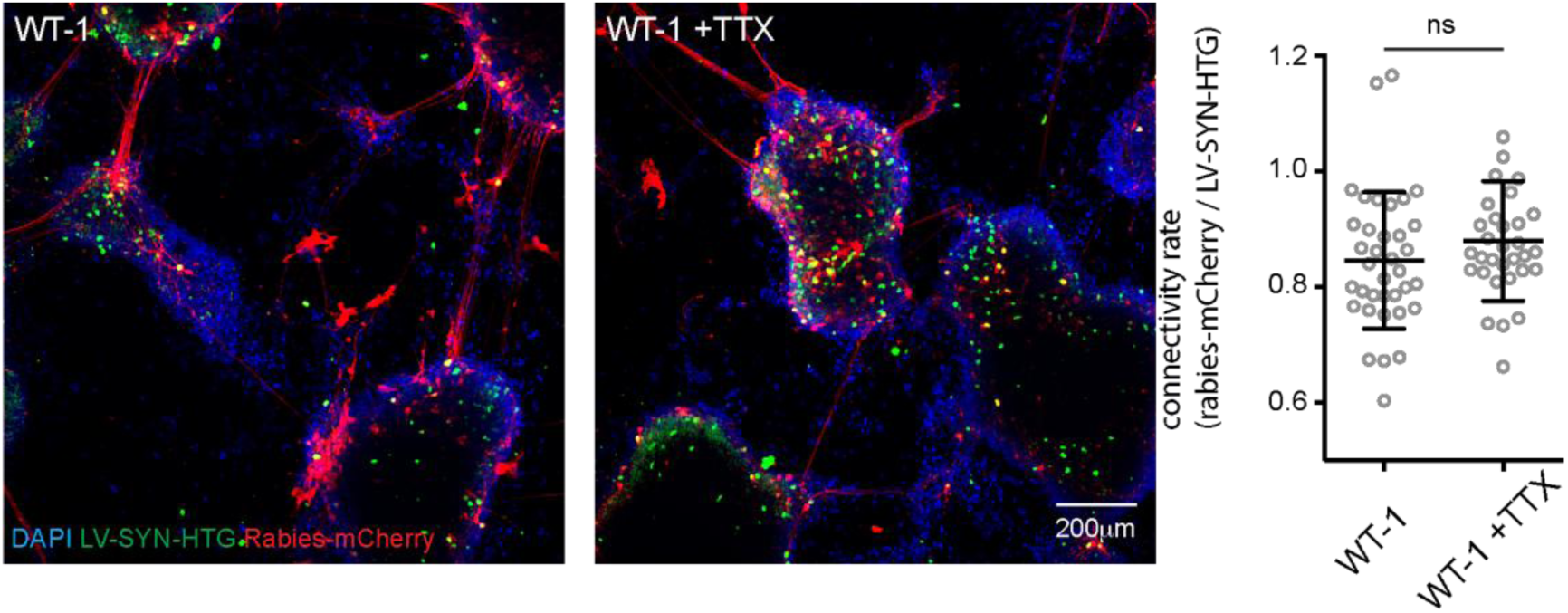
Pseudotyped rabies spread is independent of neuronal activity. Representative images of mock and TTX treated WT-1 neurons co-infected with a lentivirus expressing TVA and nuclear GFP (TVA-NLS-GFP) and a pseudotyped mCherry expressing rabies virus (Rabies-mCherry) (left). Quantification of connectivity (right). Note no difference in connectivity levels in TTX treated neuronal cultures (WT-1: 0.85 ± 0.02, n = 36 ROIs, WT-1 + TTX: 0.88 ± 0.02, n = 32 ROIs from one neuronal differentiation, one-way ANOVA.

**Figure 5—figure supplement 1.**
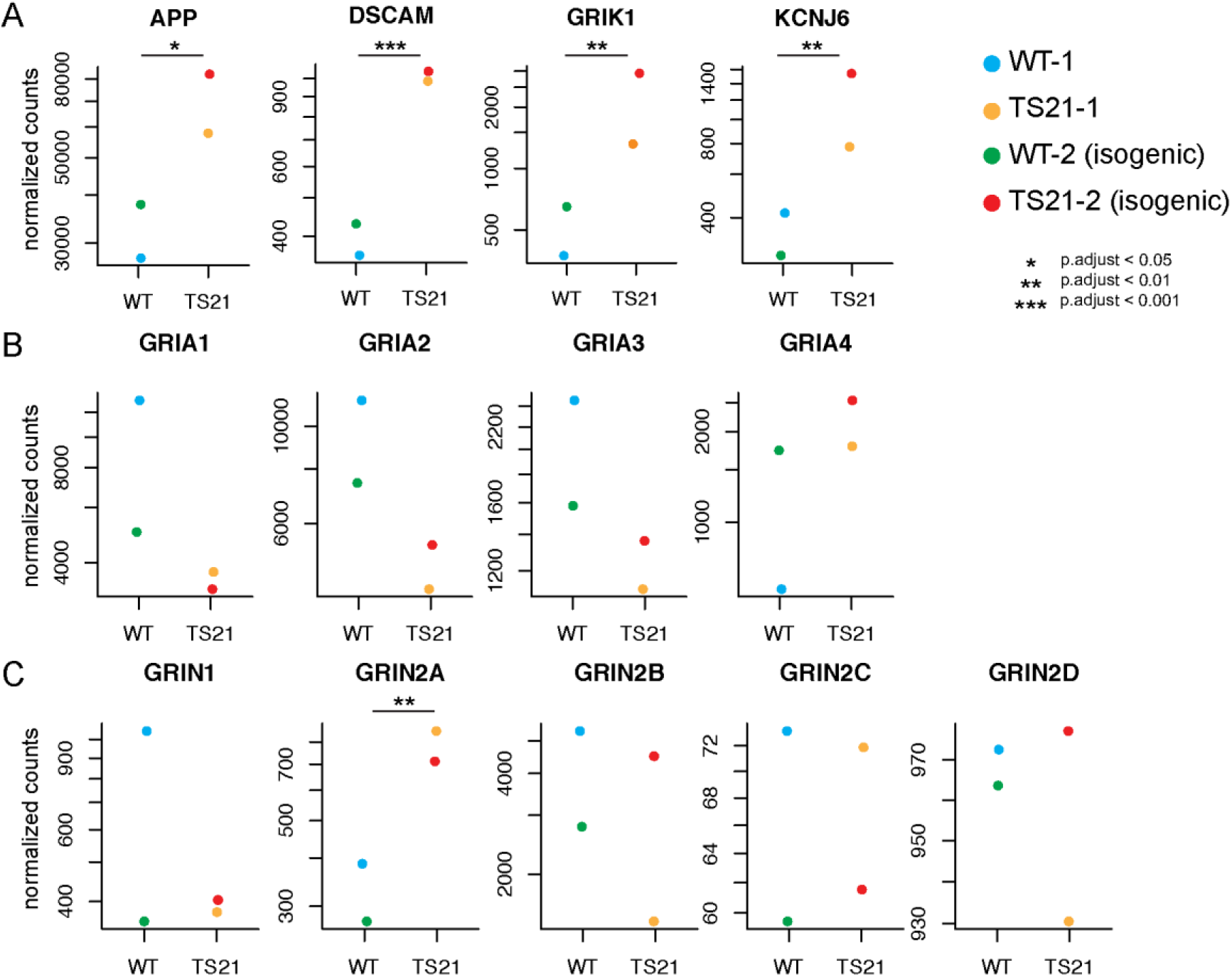
Gene expression levels of significantly up or down-regulated genes in TS21 neurons at day 50. A: genes located on chromosome 21 are significantly up-regulated in TS21 neurons. B: Expression of AMPA receptor subunits, B: Expression of NMDA receptor subunits (n =1 sample from one neuronal induction per genotype).

**Figure 5—figure supplement 2.**
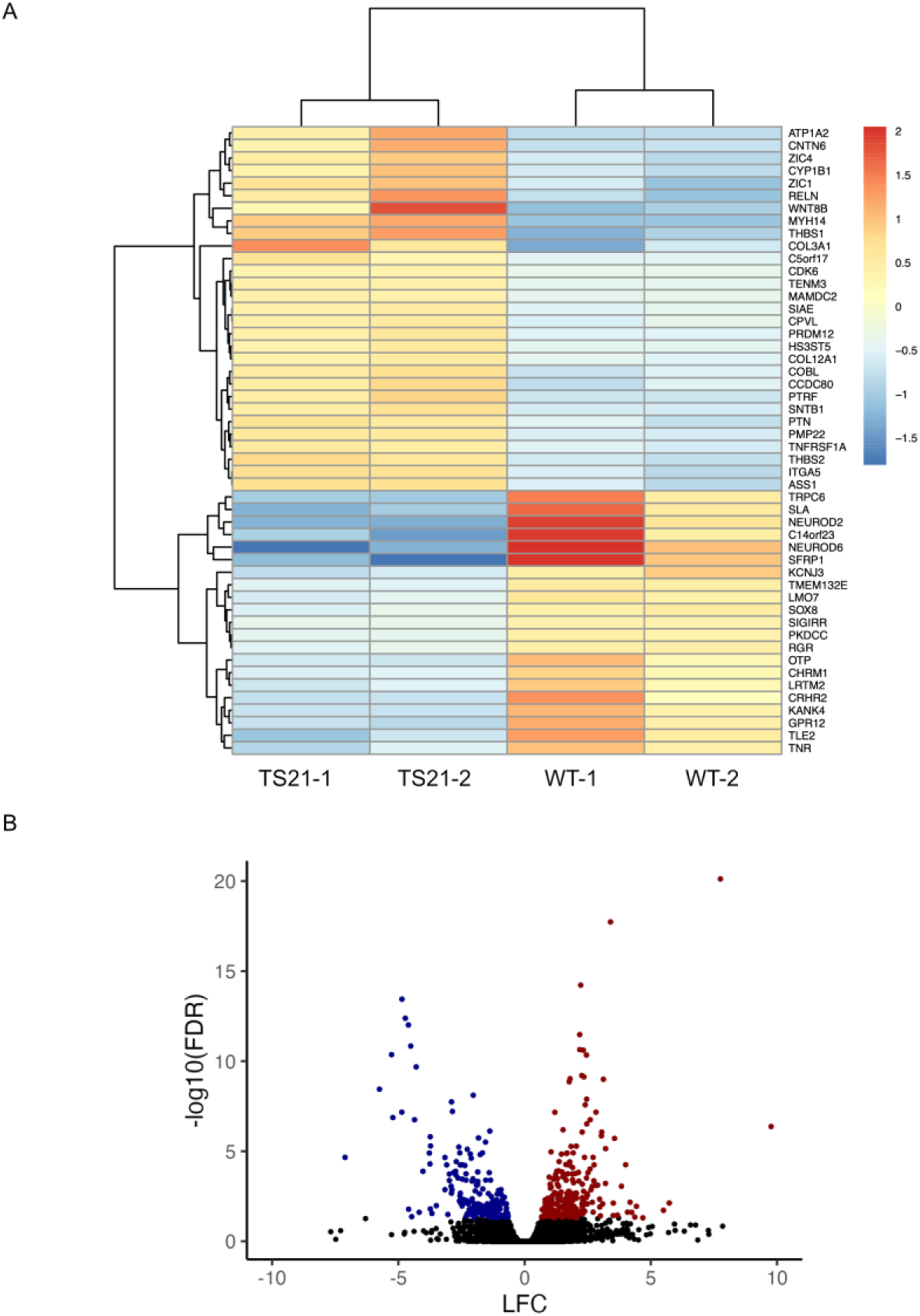
RNAseq analysis of day 50 cultures. A: Heatmap of top 50 differentially expressed genes between WT and TS21 cultures. B: Vulcano plot of up and downregulated genes between WT and TS21 cultures. A full list of all up and downregulated genes as well as the gene ontology (GO) results is available in the Supplemental files (Suppl. Table 1).

**Figure 5—figure supplement 3.**
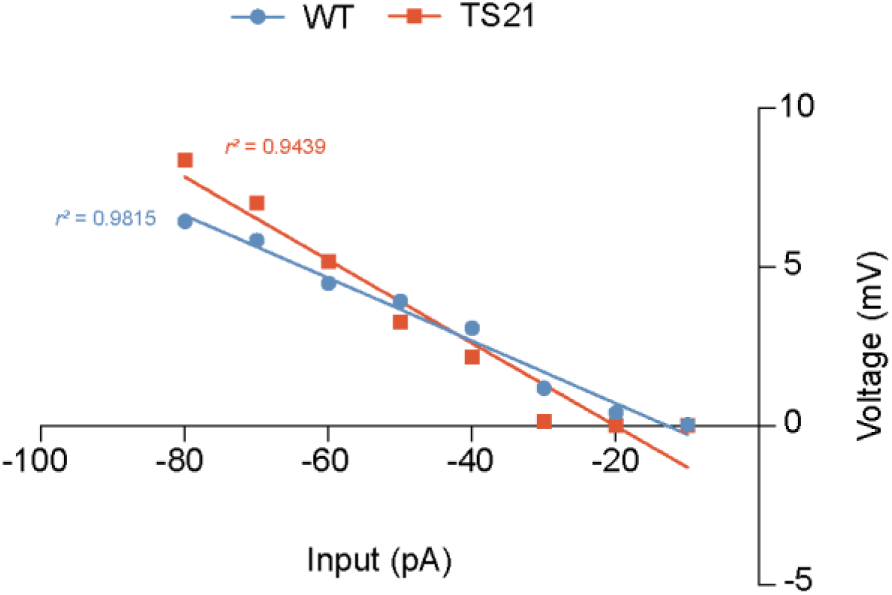
IV curves after hyperpolarizing current injection in TS21 and control neurons ex *vivo*. Positive correlation between hyperpolarization (input) and current (voltage) in WT (r2 = 0.9815) and TS21 (r2 = 0.9439) neurons ex *vivo*.

**Figure 5—figure supplement 4.**
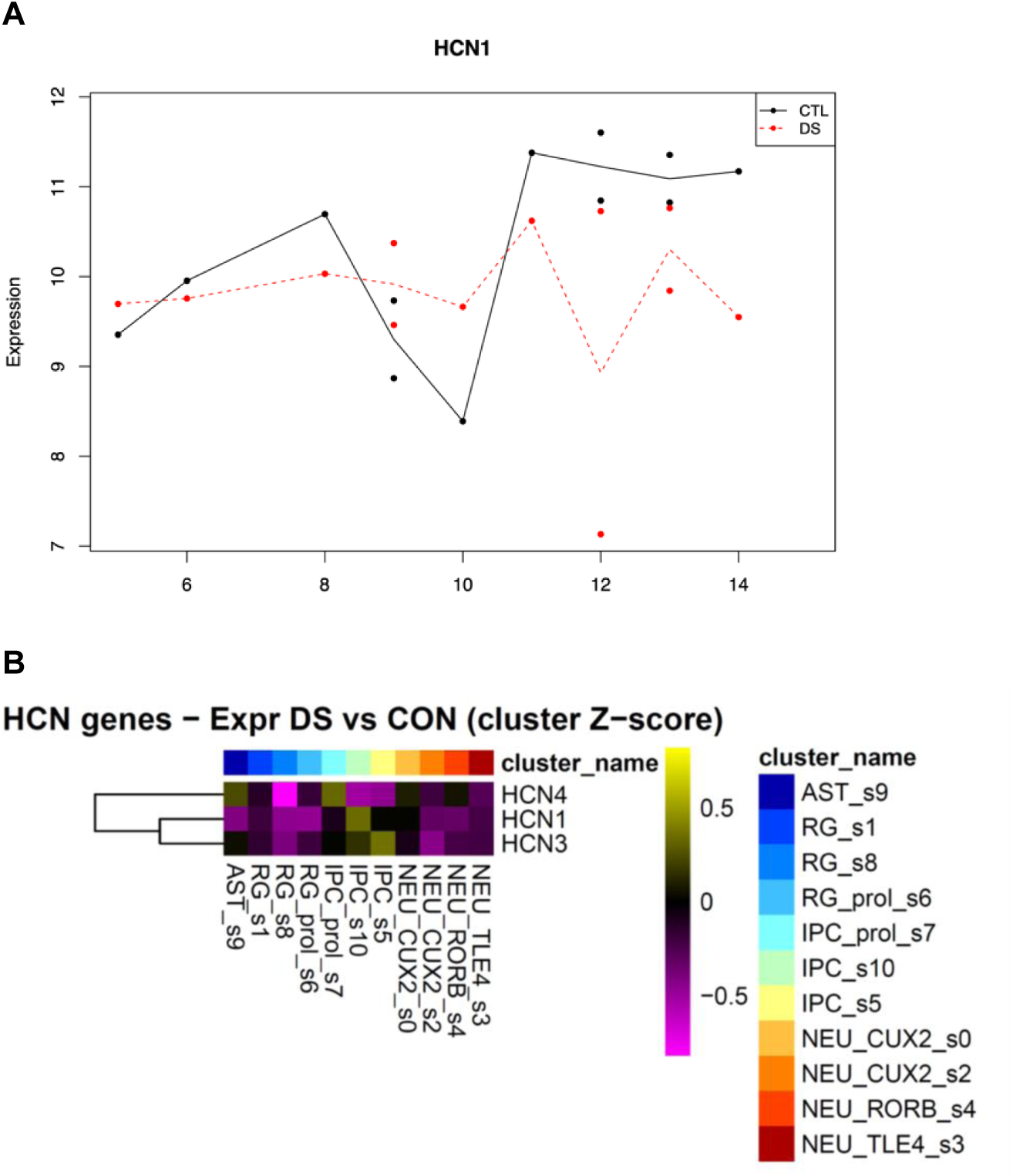
A: Expression of HCN1 in human fetal cortex and developmental period showing decrease trends in late gestation (6-8) and adult periods (12-14) (From (Olmos-Serrano et al 2016)). B: Mean expression Z-score by cluster in human fetal cortex at midgestation (N = 15 Control and 15 Trisomy 21 cases). Cortical excitatory neuron clusters are NEU_CUX2, NEU_RORB and NEU_TLE4 from the multiome data set of (Lattke et al 2026).

