## Supplemental Table 1 for "Trisomy 21 Impairs Synchronized Activity in Human Down Syndrome Cortical Excitatory Neuron Networks" (from another version of this preprint: version 1, 2024-12-27)

### Chemical synaptic transmission

| Ensembl ID | Gene Symbol | Gene name |
|---|---|---|
| ENSG00000169621 | APLF | Aprataxin and PNKP like factor |
| ENSG00000198576 | ARC | Activity regulated cytoskeleton associated protein |
| ENSG00000164326 | CARTPT | CART prepropeptide |
| ENSG00000168539 | CHRM1 | CHOLINERGIC RECEPTOR, MUSCARINIC |
| ENSG00000145920 | CPLX2 | Complexin 2 |
| ENSG00000100884 | CPNE6 | Copine 6 |
| ENSG00000170579 | DLGAP1 | DLG associated protein 1 |
| ENSG00000272636 | DOC2B | Double C2 domain beta |
| ENSG00000145864 | GABRB2 | Gamma-aminobutyric acid type A receptor beta2 subunit |
| ENSG00000177807 | KCNJ10 | Potassium voltage-gated channel subfamily J member 10 |
| ENSG00000105642 | KCNN1 | Potassium calcium-activated channel subfamily N member 1 |
| ENSG00000185760 | KCNQ5 | Potassium voltage-gated channel subfamily Q member 5 |
| ENSG00000061337 | LZTS1 | Leucine zipper tumor suppressor 1 |
| ENSG00000171532 | NEUROD2 | Neuronal differentiation 2 |
| ENSG00000112333 | NR2E1 | Nuclear receptor subfamily 2 group E member 1 |
| ENSG00000109991 | P2RX3 | Purinergic receptor P2X 3 |
| ENSG00000181195 | PENK | Proenkephalin |
| ENSG00000058335 | RASGRF1 | Ras protein specific guanine nucleotide releasing factor 1 |
| ENSG00000169071 | ROR2 | Receptor tyrosine kinase like orphan receptor 2 |
| ENSG00000063015 | SEZ6 | Seizure related 6 homolog |
| ENSG00000104888 | SLC17A7 | Solute carrier family 17 member 7 |
| ENSG00000074317 | SNCB | Synuclein beta |
| ENSG00000157005 | SST | Somatostatin |
| ENSG00000134207 | SYT6 | Synaptotagmin 6 |
| ENSG00000011347 | SYT7 | SYNAPTOTAGMIN 7 |
| ENSG00000102362 | SYTL4 | Synaptotagmin like 4 |
| ENSG00000116147 | TNR | Tenascin R |

### Voltage-gated potassium channel activity

| Ensembl ID | Gene Symbol | Gene name |
|---|---|---|
| ENSG00000164588 | HCN1 | Hyperpolarization activated cyclic nucleotide gated potassium channel 1 |
| ENSG00000182255 | KCNA4 | Potassium voltage-gated channel subfamily A member 4 |
| ENSG00000135519 | KCNH3 | Potassium voltage-gated channel subfamily H member 3 |
| ENSG00000184611 | KCNH7 | Potassium voltage-gated channel subfamily H member 7 |
| ENSG00000177807 | KCNJ10 | Potassium voltage-gated channel subfamily J member 10 |
| ENSG00000162989 | KCNJ3 | Potassium voltage-gated channel subfamily J member 3 |
| ENSG00000105642 | KCNN1 | Potassium calcium-activated channel subfamily N member 1 |
| ENSG00000185760 | KCNQ5 | Potassium voltage-gated channel subfamily Q member 5 |
